# Resistance to vincristine in cancerous B-cells by disruption of p53-dependent mitotic surveillance

**DOI:** 10.1101/2023.01.19.524713

**Authors:** Anne Bruun Rovsing, Emil Aagaard Thomsen, Ian Nielsen, Thomas Wisbech Skov, Yonglun Luo, Karen Dybkær, Jacob Giehm Mikkelsen

**Affiliations:** Department of Biomedicine, Aarhus University, Aarhus, Denmark; Lars Bolund Institute of Regenerative Medicine, BGI-Qingdao, BGI-Shenzhen, Shenzhen, China; Department of Hematology, Aalborg University Hospital, Aalborg, Denmark

## Abstract

The frontline therapy R-CHOP for patients with diffuse large B-cell lymphoma (DLBCL) has remained unchanged for two decades despite numerous phase III clinical trials investigating new alternatives. Multiple large studies have uncovered genetic subtypes of DLBCL enabling a targeted approach. To further pave the way for precision oncology, we perform genome-wide CRISPR screening to uncover the cellular response to one of the components of R-CHOP, vincristine, in the DLBCL cell line SU-DHL-5. We discover important pathways and subnetworks using gene-set enrichment analysis and protein-protein interaction networks and identify genes related to mitotic spindle organization that are essential during vincristine treatment. Inhibition of KIF18A, a mediator of chromosome alignment, using the small molecule inhibitor BTB-1 causes complete cell death in a synergistic manner when administered together with vincristine. We also identify the genes *KIF18B* and *USP28* for which CRISPR/Cas9-directed knockout induces vincristine resistance across two DLBCL cell lines. Mechanistic studies show that lack of *KIF18B* or *USP28* counteracts a vincristine-induced p53 response involving the mitotic surveillance pathway (USP28-53BP1-p53). Collectively, our CRISPR screening data uncover potential drug targets and mechanisms behind vincristine resistance, which may support the development of future drug regimens.

**Key points:**

- Inhibition of the mitotic surveillance pathway (USP28-53BP1-p53) and KIF18B induces resistance to vincristine
- Substantial synergistic effects observed when using the KIF18A-inhibitor BTB-1 with vincristine in eradicating GCB-subtype DLBCL cells

## Introduction

Only around 65% of diffuse large B-cell lymphoma (DLBCL) patients are cured after treatment with the frontline regimen R-CHOP consisting of the CD20-recognizing antibody rituximab, three chemotherapeutics (cyclophosphamide, doxorubicin and vincristine), and the immuno-suppressive steroid prednisone.^1^ To improve this prognosis, several changes of the R-CHOP regimen have been tested in phase III clinical trials including dose intensification, substitution of antibodies, and the addition of novel agents such as ibrutinib, lenalidomide and bortezomib.^2^ Unfortunately, this has not led to significant improvements. Recently, the POLARIX phase III trial comparing R-CHOP to Pola-R-CHP, in which vincristine was substituted with polatuzumab vedotin, a CD79b-targeting antibody-drug conjugate, showed higher progression-free survival after 2 years.^3^ However, the trial showed no difference in overall survival, and regulatory approval of this regimen has not yet been granted. Following initiation of these clinical trials, three recent studies characterized new genetic DLBCL subtypes using among other DNA sequencing to identify recurrent genetic alterations in large DLBCL patient cohorts^4–6^. These new genetic subtypes share considerable overlap, suggesting that they are rooted in biologically meaningful distinctions valuable for designing future clinical trials.

Vincristine has been widely used in the first-line treatment of blood, brain, bone, and kidney cancers among others since the 1960s.^7^ Vincristine interferes with mitosis by binding β-tubulin, which disrupts the capacity of microtubules to separate chromosomes during metaphase. This halts proliferating cells at the spindle assembly checkpoint (SAC) until the cells resolve the mitotic spindle disruption or the prolonged SAC causes apoptosis. Cells may also exit the SAC prematurely causing chromosome missegregation, which results in aneuploid cells, after which either cell cycle arrest, apoptosis or chromosome instability is seen. Despite the many years of use and extensive research into the mechanism of action and toxic side effects, it remains uncertain why some patients do not benefit from vincristine treatment and why resistance to vincristine occurs.^8,9^

Genome-wide genetic screening using CRISPR/Cas-based technologies enables unbiased identification of genes altering cellular responses to drugs.^10–12^ Using CRISPR/Cas, it is possible to generate a heterogenous population of cells in which all protein-encoding genes are knocked-out one by one in separate cells by introducing a genome-wide library of guide RNAs (gRNAs) and Cas9 endonuclease. Treatment of this population of cells with a specific drug, e.g. vincristine, allows resistant cells to be enriched and genes affecting drug resistance or sensitivity to be identified by deep sequencing of gRNA cassettes. Using this approach, we have previously uncovered the cellular drug response to rituximab, leading to the identification of the two B-cell receptor signaling-related genes *BLNK* and *BTK* as mediators of rituximab-induced death in germinal center B-cell-like (GCB)-subtype DLBCL B-cell lines.^13^

Using CRISPR-based gene inhibition and activation screening, Palmer et al investigated cross-resistance among the individual R-CHOP components in the DLBCL cell line Pfeiffer and the leukemia-like cell line K562.^14^ Also, CRISPR knockout screening in the acute lymphoblastic leukemia (ALL) cell line REH was performed to investigate resistance across the seven chemotherapeutics used to treat ALL.^15^ This work showed that sensitization of cells towards vincristine was increased by knockout of the gene encoding the small-molecule drug transporter ABCC1 as well as genes encoding key mitotic factors. Genome-wide screens in DLBCL cell lines have also been used to explore the drug response to CC-122, a cereblon E3 ligase modulater,^16^ and tazemetostat, an EZH2-inhibitor.^17^ In addition, CRISPR screens were utilized to identify gene knockouts affecting cell survival and proliferation in DLBCL cell lines during normal cell culture.^18–20^

Here, we use genome-wide CRISPR screening to investigate the cellular response to vincristine in DLBCL cells. We perform gene-set enrichment analysis and use FDRnet to identify subnetworks among sensitizing and resistance-inducing gene knockouts. Based on our screen data and the Bliss analysis model, we identify a small-molecular inhibitor, BTB-1, that works synergistically with vincristine to eradicate cancerous B-cells. Moreover, we identify vincristine resistance genes for which knockout interferes with vincristine-induced p53-responses, demonstrating an origin to vincristine resistance in the mitotic surveillance pathway.

## Methods

### Cell lines

The DLBCL cell lines OCI-Ly7 and SU-DHL-5 were kindly provided by Jose Martinez-Climent (University of Navarra) and obtained from the German Collection of Microorganisms and Cell Cultures (DSMZ), respectively. The cells were cultured in RPMI 1640 supplemented with 10% fetal bovine serum and 1% penicillin/streptomycin. HEK293T cell line was obtained from ATCC. They were cultured in DMEM with 5% fetal bovine serum and 1% penicillin/streptomycin. All cell lines were examined for mycoplasma infection on a regular basis. OCI-Ly7 and SU-DHL-5 were authenticated by DNA barcoding using the sensitive AmpFISTR Identifiler PCR amplification kit (Applied Biosystems). The amplified products were analysed by capillary electrophoresis (Eurofins Medigenomix GmbH, Applied Genetics). The resulting FSA file was analysed using the Osiris software (ncbi.nlm.nih.gov/projects/SNP/osiris), and the identity of the cell lines was established by comparing to DNA barcoding profiles for known cell lines obtained with the same markers and made publicly available at DSMZ.

### Plasmids

lentiCas9-Blast (Addgene #52962) and lentiGuide-Puro (Addgene #52963) were kindly provided by Feng Zhang. The Brunello pooled library of gRNAs (Addgene #73178) was kindly provided by David Root. plentiGuide-Puro with MS4A1-targeting gRNA was generated as described by Thomsen et al.^13^

### Lentiviral vector production, transduction and titration

For lentiviral vector production 4×10^6^ HEK293T cells were seeded and transfected 24 hours later with 3 µg pRSV-REV, 13 µg pMDIg/pRRE, 3.75 µg pMD.2G and 13 µg of the desired lentiviral transgene vector. Transfection was performed using calcium phosphate buffers. Medium was changed 24 hours after transfection, and 48 hours after transfection viral supernatants were harvested and filtered through a sterile filter (0.45 µm). SU-DHL-5 cells were transduced by seeding 4×10^5^ cells in a 12-well plate and then adding the viral vector preparation to a total volume of 1 mL. Titration was performed 96 hours after transduction by quantification of the proviral element WPRE in genomic DNA, as described by Ryø et al.^21^

### Genome-wide CRISPR screening

The plasmid library was amplified as described by Thomsen et al.^22^ The screen was performed in a SU-DHL-5/Cas9 clone validated by generating 90% indels (R2 = 0.91) using an *MS4A1*-targeting gRNA. The Brunello library of gRNAs was delivered to the SU-DHL-5/Cas9 clone in duplicates using lentiviral vectors with a transductional titer of 1.3×10^6^ IU/mL. To obtain an MOI of 0.5 with 77,441 gRNAs being represented in the screen on average 1,000 times, 1.55×10^8^ cells were transduced per replicate. Two days after transduction, puromycin selection was initiated and maintained for one week. Ten days after transduction, 2×10^8^ cells were harvested for genomic DNA extraction, whereas the remaining cells were resuspended in fresh medium with either saline or vincristine added. On day 31 after transduction (3 weeks of drug selection), a minimum of 1×10^8^ cells were harvested for genomic DNA extraction using NaCl precipitation, as described by Thomsen et al.^22^ PCR amplification of the gRNA cassette and subsequent gel extractions were performed as described by Thomsen et al.^22^

### Next generation sequencing

Next-generation sequencing was carried out at BGI-Research, Shenzhen. Briefly, PCR amplicons were processed by end repair and ligated to BGISEQ-500 sequencer-compatible adapters. DNA nanoball (DNB)-based sequencing libraries were generated by rolling circle amplifications. The quality and quantity of the sequencing libraries were assessed using Agilent 2100 BioAnalyzer (Agilent Technologies). Finally, the DNB libraries were sequenced with the BGISEQ-500 sequencer (MGI Tech.) with 100 paired-end sequencing with 7.7×10^7^ reads per sample replicate on average.

### Data analysis of genome-wide CRISPR screening

Cutadapt 2.10 was used to trim the NGS reads to only include the gRNA sequence by detailing the set sequence before and after the gRNA sequence. An error rate of 0.3 and a read length between 17 and 23 were allowed. Bowtie 1.3.0 was used to map the trimmed reads to the Brunello library using the -v 0 -m 1 alignment setting. The gene sets of core essential genes and non-essential genes were from Hart et al.^23^ AUC values of gene sets were calculated using the script Calc_auc_v1.1.py from https://github.com/mhegde/auc-calculation. The gene scores were calculated using the algorithm JACKS developed by Allen et al (https://github.com/felicityallen/JACKS).^24^

Gene set enrichment analysis was performed using ShinyGO 0.76.3 (http://bioinformatics.sdstate.edu/go/).^25^ The FDR cutoff was set to 0.05 and the pathway size set to min. 2 and max. 2000. The 267 and 166 genes with the highest and lowest JACKS score, respectively, and a p-value below 0.01, common for both high- and-low dose vincristine samples, were analysed.

To identify the most significant subnetworks, the algorithm FDRnet developed by Yang et al^26^ (https://github.com/yangle293/FDRnet) was used with the STRING protein-protein interaction network version 11.5 and FDR values calculated using Benjamini-Hochberg’s method from P-values calculated using JACKS gene scores. The STRING protein-protein interaction network was filtered to only include interactions with high confidence. This was done by only including interactions with a STRING-reported ‘combined score’ above 500 except for genes with less than 5 interactions where all interactions were included to ensure that all genes were represented. This resulted in a database of 384,290 protein-protein interactions. The software tool Cytoscape version 3.8.2 with the packages DyNet and clusterMaker was used to visualize the identified subnetworks. Plots were generated in R using the packages ggplot2, ggrepel, and ggprism and modified in Adobe Illustrator version 2020.

### Generation of gene knockout in cells by nucleofection, indel analysis and deep sequencing

3.2 µg chemically modified (2’-O-Methyl at 3 first and last bases, 3’ phosphorothioate bonds between first 3 and last 2 bases) gRNA (Synthego) and 6 µg Cas9 protein (Alt-R S.p. Cas9 Nuclease V3, Integrated DNA Technologies) were incubated at 25°C for 15 minutes before mixing with 6×10^5^ cells, which were rinsed and suspended in 20 µL OptiMEM. The cells were nucleofected using a 4D-nucleofector device (Lonza) in 20-µL Nucleocuvette strips (Lonza). Cells were nucleofected using the program CM-125 or CM-130 set to P3 buffer. Following nucleofection, the cells were seeded in 100 µL pre-heated medium in a 96-well plate. The following day, the cells were transferred to a 48-well plate and 100 µL medium was added before the cells were expanded.

Indel analysis was performed as described by Ryø et al^21^ using ICE software tool (Synthego, https://ice.synthego.com/). Deep sequencing of cut-site was done by Eurofins using INVIEW CRISPR CHECK and 2^nd^ PCR Protocol. CRISPResso2 was used to analyze the raw NGS reads for genome editing (http://crispresso2.pinellolab.org/).^27^

### Flow cytometry-based assays

Prior to the seeding of cells, the proliferation rate was aligned by passing the cells every second day and seeding them at the same density at either 4×10^5^ or 5×10^5^ cells/ml for SU-DHL-5 and OCI-Ly7, respectively. This was done a minimum of 3 passages before the assay initiation. 7.2×10^4^ SU-DHL-5 or 9.0×10^4^ OCI-Ly7 cells were seeded in a 96-well plate in 175 µL medium in a random set-up to avoid bias from the flow cytometer running order.

In drug assays, drug, DMSO, or saline was added in 5 µL. After 48 to 60 hours, the cells were passaged by removing and replenishing 120 or 100 µL medium for SU-DHL-5 or OCI-Ly7 cells, respectively. In drug assays comparing gene knockouts to wildtype, vincristine or saline was added as well when passaging the cells, whereas only medium was replenished in drug synergy assays. After 110-120 hours, the samples were mixed and 97 µL cell suspension was transferred to a V-bottom 96-well plate with 2.5 µL 50µg/mL propidium iodide. The number of live and dead cells in 30 µL was counted using a NovoCyte 2100YB Flow Cytometer (Agilent Technologies). Vincristine sulfate (Merck) was dissolved and diluted in saline. BTB-1 (Axon Medchem) was dissolved in DMSO and diluted in saline.

For intracellular staining of p53, cells from 3 wells were pooled and washed in 1% BSA (bovine serum albumin) in PBS and then stained with Fixable Viability Dye eFluor 780 (Invitrogen) as viability marker for 15 min. at 4°C. Cells were washed again before using ice-cold methanol for fixation and permeabilization for 2 min, and then washed and stained with PE-conjugated p53 antibody (Clone DO-7, BioLegend, Cat. 645805) 1:100 in 0.1% Triton X-100 0.5% BSA in PBS at room temperature for 30 min prior to a wash in 1% BSA in PBS, and run on NovoCyte Quanteon 4025 Flow Cytometer (Agilent Technologies).

For the cell cycle assay, cells from 3 wells were pooled and the 488 EdU Click Proliferation Kit (BD Biosciences) was used using Fixable Viability Dye eFluor 780 (Invitrogen) as viability marker and ice-cold methanol for fixation and permeabilization. For the apoptosis assay, the CellEvent Caspase-3/7 Green Detection kit (Invitrogen) was used with propidium iodide as a viability marker. Nutlin-3 (Merck) was dissolved in DMSO and diluted in saline.

### Western blotting

1×10^6^ cells were washed in PBS and lysed in RIPA Lysis and Extraction buffer (ThermoFisher Scientific, Cat. 89901) supplemented with 10 mM NaF and 1x complete protease inhibitor cocktail (Roche). Then, XT Sample Buffer, 4x (Biorad, Cat. 161-0791) and XT Reduction Detergent 20x (Biorad, Cat. 161-0792) were added to the samples, and the samples were boiled for 5 minutes before being loaded on the gel (4-15% Criterion TGX Precast Midi Protein Gel, Bio-Rad). Proteins were separated by SDS-PAGE and blotted into a polyvinylidene fluoride (PVDF) membrane. Membranes were divided in two by cutting at the 100 kDa mark. Membranes were blocked in 5% skim milk in TBS supplemented with 0.005 % Tween-20 (Sigma-Aldrich). The bottom and top parts of the membrane were then incubated overnight at 4°C with primary p53 antibody (2 µg/mL, 1:250) (Clone DO-1, BD Biosciences, Cat. 554293) and primary vinculin antibody (1:10,000) (Clone hVIN-1, Sigma Aldrich Cat. V9131), respectively. The membranes were then washed in TBS supplemented with 0.005 % Tween-20 (Sigma-Aldrich) and incubated with horseradish peroxidase (HPR)-conjugated secondary antibody and visualized by chemiluminescence using a horseradish peroxidase substrate (Bio-Rad, Cat. 170-5061). Quantification was performed using ImageJ.

### RNA extraction and qPCR of gene expression

1×10^6^ cells were lysed with 0.5 mL TRI ReagentTM Solution (Fischer Scientific, Cat. AM9738), followed by the addition of 0.1 mL Chloroform. After mixing and 3 minutes of incubation at room temperature, the samples were centrifuged at 17.000g for 15 minutes at 4℃. Hereafter, 300 µL of the aqueous phase was mixed with 375 µL ethanol and loaded onto a column from the hiPure miRNA purification kit (Roche, Cat. 5080576001). The samples were then washed and eluted according to the manufacturer’s instructions.

800 ng RNA was treated with DNase I (Thermo Fisher Scientific, Cat. EN0521) in a total volume of 10 µL, for 30 minutes at 37℃. Hereafter, the DNase was inactivated by the addition of 1µL 50 mM EDTA followed by incubation at 65℃ for 10 minutes. 500 ng DNase-treated RNA was then used for cDNA synthesis in a 10 µ L reaction using Maxima H Minus cDNA Synthesis Master Mix (Thermo Fisher Scientific, Cat. M1662). 4.8 µL 10-fold diluted cDNA was subsequently used in a 10 µL qPCR reaction with 500 nM forward and reverse primer, using RealQ Plus 2x Master Mix Green (Ampliquon, Cat. A323402). The primers were designed to span an exon-exon junction to avoid contamination from genomic DNA. The qPCR was run on a Lightcycler 480 II (Roche) in technical triplicates. The relative expression levels were determined using a standard delta-delta CT calculation normalized to GADPH.

### Statistical analysis

One-way ANOVA was performed in the software program GraphPad Prism version 9. If a p-value below 0.05 was calculated, then Dunnett’s multiple comparison test was used to compare the mean of the samples with the corresponding control sample.

### Data sharing statement

NGS fastq files from genome-wide CRISPR screening are available from the corresponding author upon request. Read count table and JACKS gene score table for CRISPR screening data can be found in supplementary tables.

## Results

### Uncovering genes affecting the response to vincristine by genome-wide CRISPR screening in SU-DHL-5 cells

To identify genes affecting the response to vincristine in DLBCL cells, we performed genome-wide CRISPR screening in the GCB-subtype SU-DHL-5 cell line (Cas9-expressing clone), which is more sensitive to vincristine than other DLBCL cell lines.^28^ We used the Brunello library developed by Doench et al targeting 19,114 genes with 4 gRNAs per gene on average (Figure 1A).^29^ After transduction and a selection for gRNA-containing cells only, we challenged the cells with vincristine at a low (1.3 ng/mL) and high (1.7 ng/mL) dosage (table 1) in two independent replicates for 3 weeks before harvesting cell pellets for NGS sequencing.

**Table 1.** Vincristine dose response values from Falgreen et al.^30^ GI50: 50% cell growth inhibition. TGI: total growth inhibition. LC50: 50% cell death.

| Cell line | GI50, ng/mL | TGI, ng/mL | LC50, ng/mL | Cell seeding, cells/mL |
| --- | --- | --- | --- | --- |
| SU-DHL-5 | 0.71 | 1.00 | 1.70 | 2.5x10<sup>5</sup> |
| OCI-Ly7 | 2.55 | 2.90 | 3.39 | 2.5x10<sup>5</sup> |

**Figure 1.**
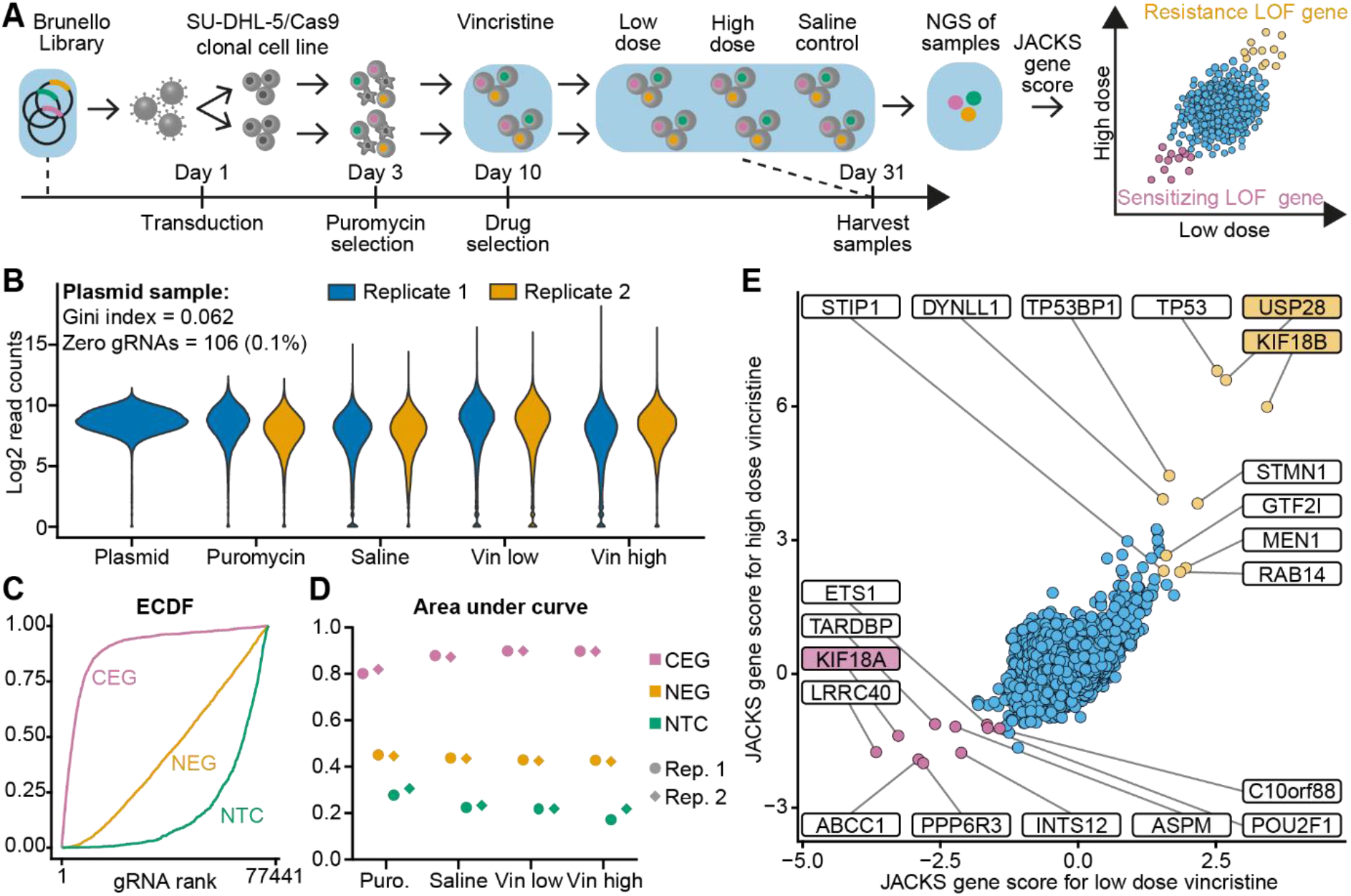
Genome-wide CRISPR screening using vincristine as selective pressure. (**A**) The genome-wide CRISPR screening process and how the samples indicated with blue background were generated. (**B**) Distribution of log2-normalized read counts of gRNA sequences in NGS data. (**C**) Empirical cumulative distribution functions (ECDF) plotted for gRNAs targeting core essential genes (CEG), non-essential genes (NEG), and non-targeting controls (NTC). Optimally, gRNAs targeting CEG should have a low rank, due to successful gene knockout causing cell death. (**D**) Area under the curve (AUC) values plotted for ECDFs across the three gene sets and all samples. (**E**) JACKS gene scores for the low- and high-dose vincristine samples plotted against each other.

The coverage of each gRNA in each sample was on average 450 and 323 at minimum. The plasmid sample had a Gini index^31^ of 0.062, and less than 0.1% of the gRNAs were unaccounted for (Figure 1B). To ensure high screen quality, we used the core essential and non-essential gene sets established by Hart et al^23^ and made empirical cumulative distribution functions (ECDF) of gRNA rankings (Figure 1C). All samples showed high area under curve (AUC) values of the ECDF near or above 0.90 for the core essential genes (Figure 1D). This demonstrated a high efficiency in generating gene knockout and selecting cells based on the phenotypic consequence across all samples.

We compiled the gRNA read counts into gene scores using the JACKS algorithm, which takes varying gRNA efficiency into account when assigning gene scores.^24^ Saline-treated replicates were used as baseline against the two replicates for each vincristine concentration, resulting in JACKS gene scores for high and low vincristine dosages. Genes for which Cas9-induced loss-of-function (LOF) made the cells more resistant towards vincristine were defined by high JACKS scores, whereas low JACKS scores pointed to LOF genes rendering the cells more sensitive to vincristine. The JACKS gene scores showed high correlation when plotting the high-dose scores against the low-dose scores, demonstrating that the same LOF genes had the highest impact across both drug dosages (Figure 1E). Notably, sensitizing LOF genes had a higher impact in the low-dose condition, whereas resistance LOF genes had a higher impact in the high-dose condition. The *ABCC1* (*MRP1*) gene encoding an ABC transporter known for mediating multidrug resistance was one of the genes with the lowest JACKS score.^32^ This served as an initial validating proof that the screening approach uncovered genes with a high impact on the cellular vincristine response.

### Significant pathways, subnetworks and protein complexes in the cellular response to vincristine

To identify more general tendencies at the pathway level, we used ShinyGO to perform and visualize gene-set enrichment analysis using Gene Ontology (GO) and Reactome databases.^25^ For the 166 genes with the lowest JACKS score and a p-value below 0.01 for both the high- and low-dose samples, gene sets related to microtubule depolymerization, mitotic spindle organization, INO80 complex, and DNA damage recognition in Global-Genome Nucleotide Excision Repair (GG-NER) were among the enriched gene sets (Supplementary Figure S1A-C). For the 267 genes with the highest JACKS score and a p-value below 0.01 for both the high- and low-dose samples, gene sets related to negative regulation of mTOR signaling, GATOR1-complex, CD40 receptor complex, and Rab GTPases regulation of trafficking were among the enriched gene sets (Supplementary Figure S2A-C).

The algorithm FDRnet was developed by Yang and colleagues to identify subnetworks of cancer drivers in genomic datasets with high scores and high connectivity within larger networks.^26^ Here, we used FDRnet to identify subnetworks and protein complexes with significant influence on the vincristine response as suggested by our screen data (Figure 2A). We used JACKS-derived FDR values and the STRING protein-protein interaction network, but only included high-confidence interactions with direct evidence of physical protein-protein interactions. Among the most sensitizing LOF genes, we identified subnetworks related to mitotic spindle organization, the cytoplasmic dynein complex responsible for trafficking along microtubules, and the INO80 complex, an ATP-dependent chromatin remodeler (Figure 2B).^33^ The subnetworks identified for resistance LOF genes included B-cell receptor (BCR) signaling, tumor necrosis factor receptor (TNFR) signaling entailing CD40 receptor signaling, membrane trafficking represented by Rab GTPases and the GATOR1 and TSC1/2 complex, which both negatively regulate mTOR signaling (Figure 2C). Except for subnetworks related to GG-NER, the FDRnet approach identified subnetworks related to the same biological processes as the most enriched gene-sets in the gene-set enrichment analysis. In addition, *TP53* was discovered to be a key gene as a central connection for many of the identified subnetworks.

**Figure 2.**
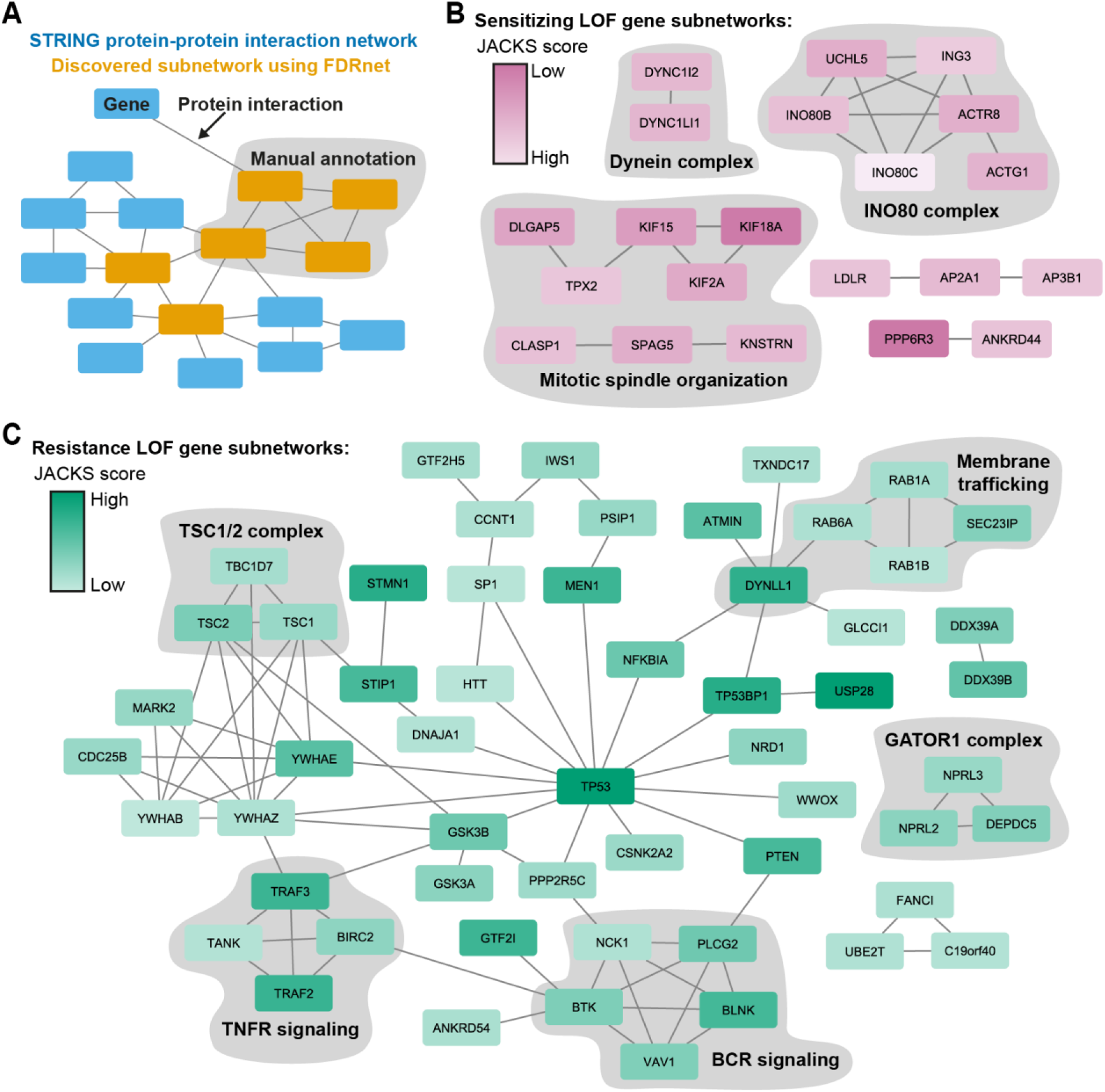
Subnetworks of important genes in the response to vincristine identified using FDRnet. (**A**) The STRING protein-protein interaction network in blue with an FDRnet-discovered subnetwork in orange. The grey clouds indicate manually annotated descriptions of the unifying biological process or protein complex. (**B-C**) The FDRnet-identified subnetworks among the most sensitizing (**B**) and resistance-inducing (**C**) LOF genes.

We have previously shown that knockout of genes related to BCR signaling, including *BLNK* and *BTK*, renders GCB-subtype DLBCL cell lines OCI-Ly7 and SU-DHL-5 more tolerant towards rituximab.^13^ Notably, median-normalized read counts showed that gRNAs targeting *BLNK* and *BTK* were enriched in the saline-treated sample compared to the baseline in the plasmid sample. This implies that knockout of these genes confers increased survival signaling and proliferation during normal cell culture (Supplementary Figure S3A-B). To identify the subnetworks that impacted the survival and proliferation rate without vincristine, we color-coded the genes according to their JACKS scores for saline-treated samples, where the initial plasmid sample was used as baseline (Supplementary Figure S4A-C). Besides BCR signaling, genes related to TNFR signaling and the GATOR1 complex and the genes *PTEN* and *NFKBIA* also had high JACKS scores for the saline-treated samples. Notably, all these subnetworks and genes are involved in NF-κB and PI3K-AKT-mTOR signaling, which regulates survival signaling, proliferation, and stress response.^34,35^ This indicates that increased tolerance to vincristine can be induced by LOF genes leading to more proliferation and survival signaling in general.

In summary, LOF genes related to mitotic spindle organization sensitize the cells to vincristine with little impact during normal cell culture, and *TP53* is a central component connecting the subnetworks that increase the tolerance to vincristine.

### KIF18A-inhibitor BTB-1 acts synergistically with vincristine in eradicating DLBCL cells

Our screen uncovered genes related to mitotic spindle organization to be essential for the DLBCL cells to survive vincristine treatment. Median-normalized read counts for gRNAs targeting *KIF18A*, a kinesin-8 motor family member involved in chromosome alignment during mitosis,^36^ showed that gene knockout sensitized the DLBCL cells to vincristine without affecting the saline-treated samples (Figure 3A). We used the KIF18A-inhibitor BTB-1^37^ to test if the screening results could be used to identify targets for small-molecule inhibitors that act synergistically with vincristine in killing DLBCL cells.

**Figure 3.**
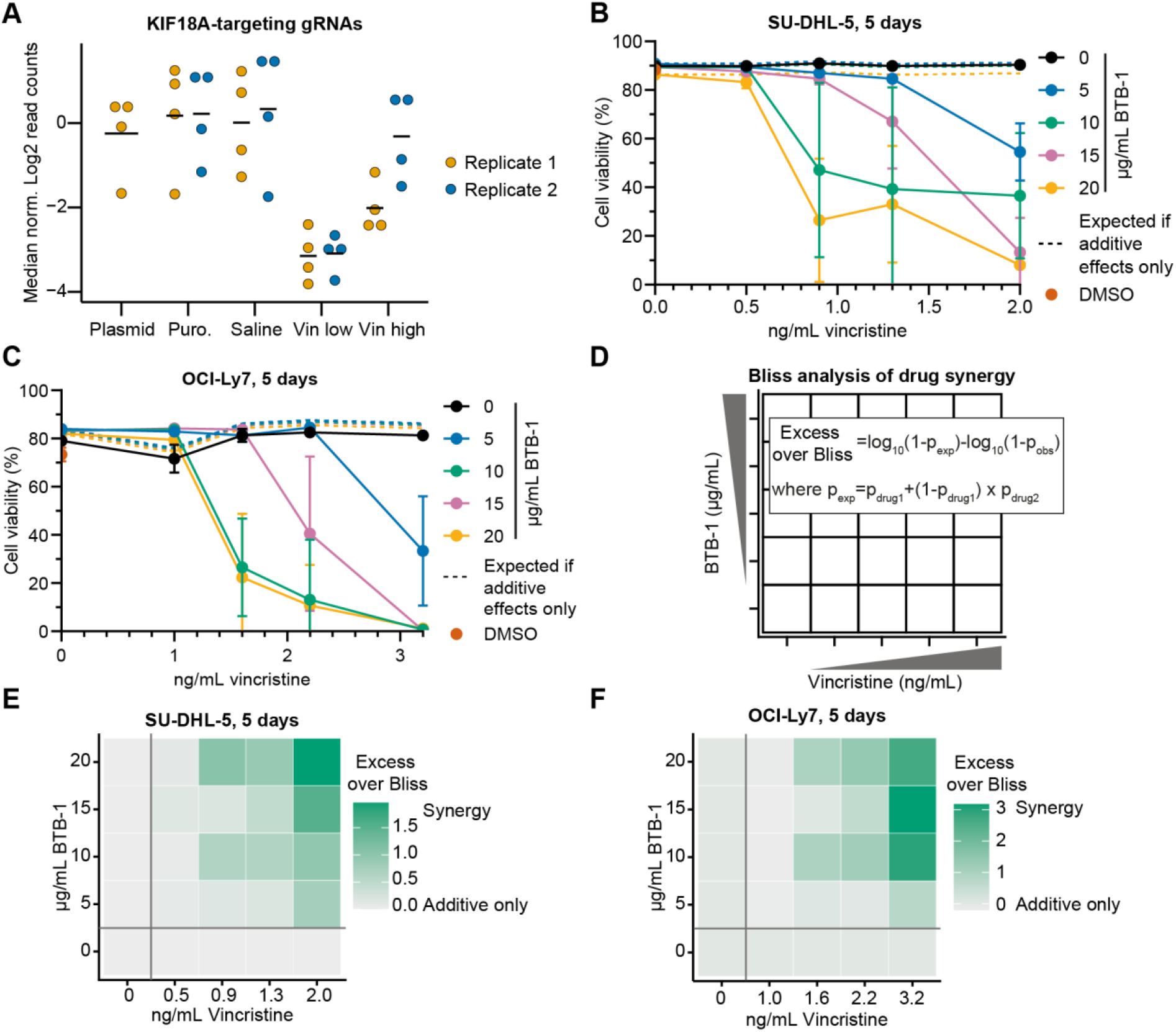
The KIF18A-inhibitor BTB-1 combined with vincristine shows synergistic effect in eradicating DLBCL cells. (**A**) Median normalized log2 read counts of gRNAs targeting *KIF18A* in the genome-wide CRISPR screening. (**B-C**) Drug assays in SU-DHL-5 (**B**) and OCI-Ly7 (**C**) cells showing the synergistic effect of vincristine and BTB-1. The dotted lines indicate theoretically calculated values showing the expected effect if the individual drug effect combined was only additive. These values were calculated using the Bliss analysis model. Drug assays were performed by counting live and dead cells after propidium iodide viability staining on a flow cytometer. (**D**) The Bliss analysis used to model the individual drug effects to test whether the drug interaction effect is synergistic or additive. Here p_x_ denotes the proportion of cells at the particular dose of the particular drug(s) compared to saline-treated cells. p_drug1_, p_drug2_, and p_obs_ is the proportion for vincristine, BTB-1, and combined, respectively. (**E-F**) The Bliss analysis for cell counts of live cells for SU-DHL-5 (**E**) and OCI-Ly7 (**F**) cells.

Treating SU-DHL-5 cells with either vincristine (dosages ranging from 0 to 2.0 ng/mL) or BTB-1 (dosages from 0 to 20 µg/mL) one at a time did not result in cell death. However, the combination of the two drugs led to extensive cell death with the highest drug concentrations resulting in less than 6% viable cells (Figure 3B). A similar pattern was observed in OCI-Ly7 cells, in which combinations of vincristine (0-3.2 ng/mL) and BTB-1 (0 to 20 µg/mL) led to complete cell death, whereas the drugs individually had no measurable effect (Figure 3C). Robust synergistic effects of using BTB-1 together with vincristine was confirmed using the Bliss Independence Model^14^ (Figure 3D) on cell counts of viable cells for SU-DHL-5 (Figure 3E) and OCI-Ly7 (Figure 3F). Together, our findings demonstrate the capacity of genome-wide CRISPR screening to uncover synergistic drug interactions exemplified by the substantial synergistic effect of BTB-1 on vincristine-induced cell death.

### *KIF18B* and *USP28* knockout increases tolerance towards vincristine

Together with *TP53*, the genes *KIF18B* and *USP28* had significantly higher JACKS scores than all other genes, and gRNAs targeting these genes only showed strong enrichment in samples treated with vincristine, but not in saline-treated samples (Figure 4A-B). *KIF18B* encodes, like *KIF18A*, a kinesin-8 motor family member involved in mitotic spindle organization by regulating astral microtubules,^38^ whereas *USP28* encodes a de-ubiquitinase regulating many cancer-related pathways.^39^ To generate SU-DHL-5 and OCI-Ly7 cell lines with knockout of either *KIF18B* or *USP28*, we nucleofected cells separately with two gRNAs for each gene, generating robust indel rates around 90% and at minimum 72% after 8 days. To ensure the stability of the established knockout populations, we confirmed comparable indel rates and indel patterns 25 days or more after the initial measurement (Figure 4C and Supplementary Figure S5A-B). Cells treated with two control gRNAs targeting safe-harbor loci were also included (Supplementary Figure S5B).

**Figure 4.**
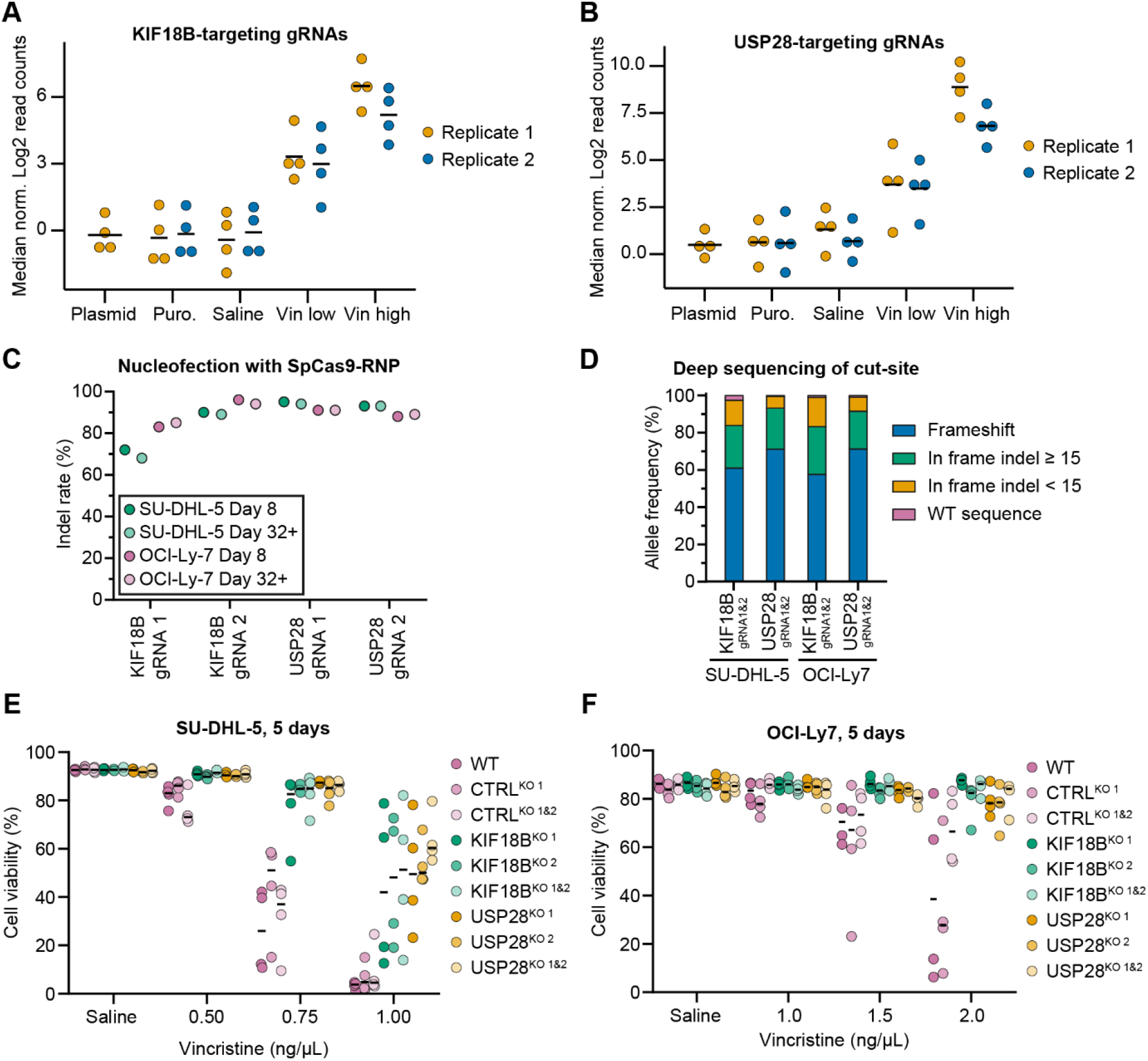
*KIF18B* and *USP28* knockout induces resistance to vincristine. (**A-B**) Median normalized log2 read counts of gRNAs targeting *KIF18B* (**A**) and *USP28* (**B**) in the genome-wide CRISPR screening. (**C**) Indel rates measured after Cas9-induced gene knockout using Sanger sequencing at day 8 or more than 32 days after nucleofection. (**D**) Allele frequencies of indels in double knockout generated cell lines by deep sequencing (>119,000 reads) of the cut-site. (**E-F**) Drug assays in SU-DHL-5 (**E**) and OCI-Ly7 (**F**) generated gene knockout cell lines. Drug assays were performed by counting live and dead cells after propidium iodide viability staining on a flow cytometer.

In addition, to further increase the knockout rate, cell populations generated using the first gRNAs were nucleofected with the second gRNAs targeting the same gene. Deep sequencing of the gRNA-targeted *KIF18B* and *USP28* loci verified extensive gene disruption (Figure 4D). Hence, less than 3% of the sequenced alleles mapped to WT sequence in SU-DHL-5 and OCI-Ly7 cells, and more than 83% of the sequenced alleles had frameshifts or indels spanning 15 or more nucleotides. For both loci, the two gRNAs recognizing closely located binding sites induced large deletions above 70 bp in more than 15% of the sequenced alleles (Supplementary figure S5C-D). The high frequency of frameshift mutations and large deletions led to reduced mRNA expression due to nonsense-mediated mRNA decay for the *USP28* transcript, but not for *KIF18B* (Supplementary Figure S5E-F). For *KIF18B*, however, the gRNAs were designed to target exon 1 encoding the essential motor domain of KIF18B protein responsible for binding microtubules.

In both SU-DHL-5 and OCI-Ly7 cells, knockout of either *KIF18B* or *USP28* strongly increased the tolerance towards vincristine. For SU-DHL-5 cells treated with 0.75 ng/mL vincristine twice during 5 days, the average viability of naïve cells and the control cells generated using irrelevant gRNAs was below 44%, whereas the *KIF18B-* and *USP28-*deficient cells were more resistant with viabilities above 76% (Figure 4E). For OCI-Ly7 cells cultured in 2.0 ng/mL vincristine, the average viability for naïve and control gRNA-generated cells was between 34% and 68%, whereas the *KIF18B-* and *USP28*-deficient cells showed viabilities comparable to the saline condition above 77% (Figure 4F). In summary, these data demonstrate that lack of either *KIF18B* or *USP28* renders GCB-subtype DLBCL cells more resistant to vincristine.

### KIF18B and USP28 mediate accumulation of activated p53 during vincristine treatment

The FDRnet analysis revealed *TP53* as a central gene connecting most subnetworks found among resistance LOF genes including a connection to *USP28* through *TP53BP1*. In fact, upon prolonged mitosis, USP28 has been shown to de-ubiquitinate and, thus, stabilize p53 through binding the scaffold protein 53BP1 (encoded by *TP53BP1*) leading to cell cycle arrest and apoptosis.^8,40^ We speculated that resistance to vincristine induced by *KIF18B* and *USP28* deficiency could be explained by their influence on restricting this p53 response.

To study the role of p53, we first examined the effect of vincristine on p53 expression. Western blotting showed an increased level of p53 upon vincristine treatment in SU-DHL-5 cells, whereas the effect on p53 was less pronounced in *USP28* knockout cells (Figure 5A-B). OCI-Ly7 cells showed a constitutively high level of p53 protein, which did not increase further upon vincristine treatment (Figure 5A-B). Notably, *TP53* in OCI-Ly7 cells carries a variant causing a glycine to aspartic acid change in the DNA binding domain of p53 (p.G245D), which destabilizes the protein and increases the amount of misfolded, nonfunctional protein.^41^

**Figure 5.**
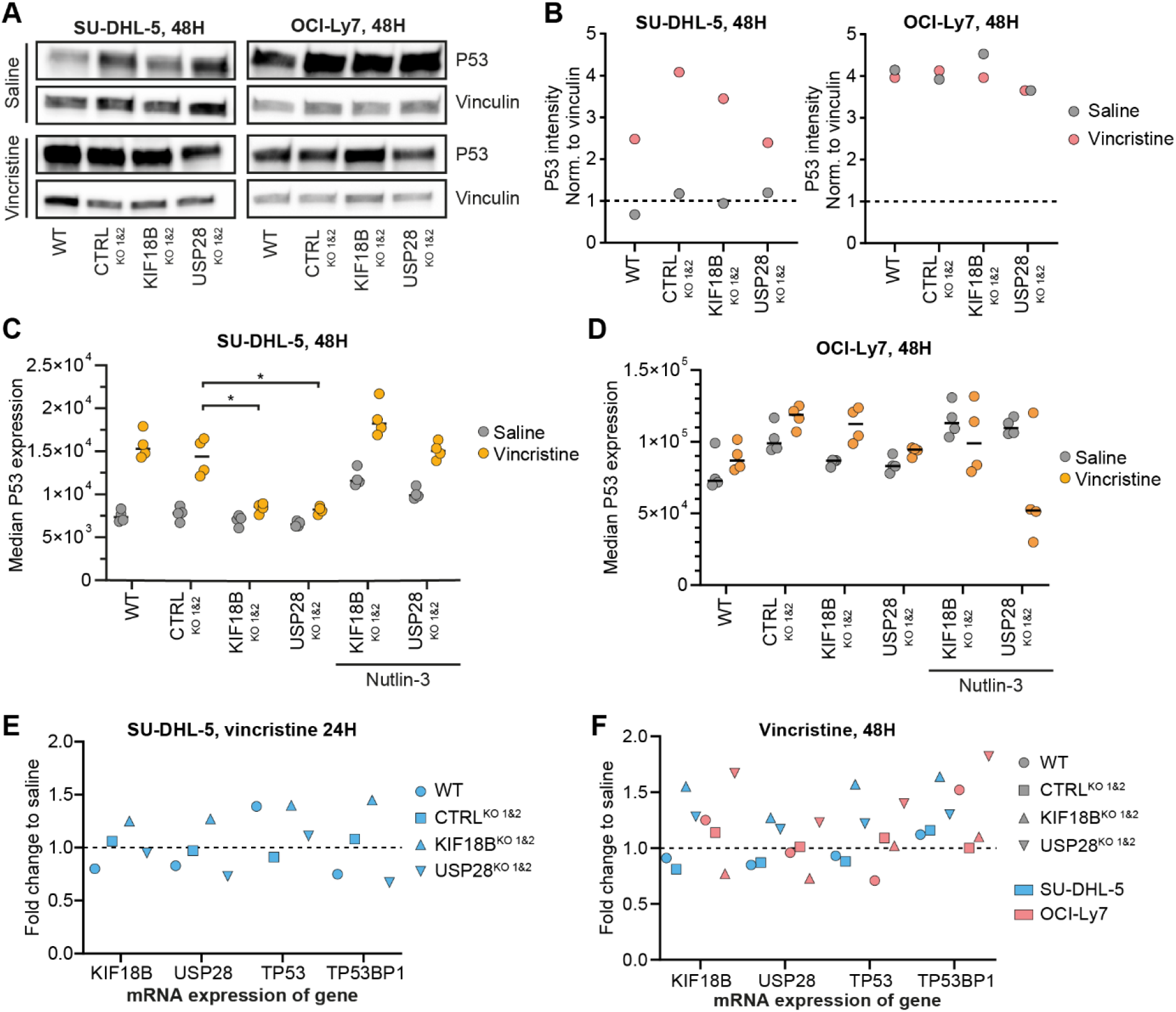
*KIF18B* and *USP28* knockout affects vincristine-induced p53 accumulation. (**A**) Western blot of p53 and vinculin protein comparing vincristine to saline treatment after 48 hours. (**B**) Quantification of Western blot performed using ImageJ. Quantification shows p53 intensity normalized to vinculin for each sample. (**C-D**) Median fluorescence intensity of intracellular staining of p53 protein determined by flow cytometry for SU-DHL-5 (**C**) and OCI-Ly7 (**D**) cells after 48 hours. (**E**) mRNA expression after 24 hours of vincristine treatment in SU-DHL-5 cells measured by qPCR. (**F**) mRNA expression after 48 hours of vincristine treatment in SU-DHL-5 and OCI-Ly7 cells measured by qPCR.

Flow cytometry-based intracellular staining of p53 showed that knockout of *KIF18B* and *USP28* in SU-DHL-5 cells drastically decreased vincristine-induced accumulation of p53 (Figure 5C). The single-cell output revealed that only some cells showed induction of p53, reflecting a dynamic nature of p53 expression (flow plots available in Appendix 2).^42^ This may explain the lack of effect of *KIF18B* knockout on p53 induction as determined by Western blotting (Figure 5A-B). p53 is negatively regulated by MDM2 by ubiquitination leading to degradation.^43^ We used nutlin-3, a small molecule inhibitor of MDM2, to inhibit MDM2 function and found in SU-DHL-5 cells that the ability of vincristine to induce p53 expression was restored in *KIF18B* and *USP28* knockout cells (Figure 5C). For OCI-Ly7, a constitutive high level of p53 protein was seen across all conditions with a median p53 level that was on average 10-fold higher than the level in SU-DHL-5 (Figure 5D). In these cells, inhibition of MDM2 using nutlin-3 increased the level of p53 in the saline-treated samples, but the effect could not be detected in the vincristine-treated samples due to toxicity. For both cell lines, noteworthy changes of *TP53, KIF18B, USP28*, or *TP53BP1* mRNA levels were not observed after 24 or 48 hours of vincristine treatment (Figure 5E-F), suggesting that vincristine treatment did not support accumulation of p53 at the transcriptional level. Collectively, our findings suggest that KIF18B and USP28 are crucial for the accumulation of p53 upon vincristine treatment in *TP53* wildtype cells.

### KIF18B and USP28 mediate induction of cell cycle arrest and apoptosis upon vincristine treatment

We speculated that reduced p53 accumulation in KIF18B- and USP28-deficient cells would lead to less P21 activation and thus, less vincristine-induced cell cycle arrest and apoptosis. Cell cycle analysis in SU-DHL-5 cells showed that vincristine treatment in control samples (naïve cells and cells treated with irrelevant gRNAs) reduced the percentage of cells actively replicating DNA in S-phase from 64% in saline-treated samples to 19% (Figure 6A). For OCI-Ly7, vincristine treatment reduced the percentage of cells in S-phase from 64% in saline-treated samples to 40% (Figure 6B). For both SU-DHL-5 and OCI-Ly7, knockout of *KIF18B* led to 10% more cells in S-phase compared to control samples. Likewise, knockout of *USP28* increased the number of cells in S-phase in SU-DHL-5 with 19%, but not for OCI-Ly7. In both SU-DHL-5 and OCI-Ly7, treatment with nutlin-3 increased the level of p53 and reverted the effect of *KIF18B* and *USP28* knockout (Figure 6A-B). Notably, treatment of saline-treated samples with nutlin-3 did only vaguely affect the induction of cell cycle arrest, suggesting that cell cycle arrest was not only affected by the p53 level, but also by the vincristine-driven activation of p53. Notably, vincristine treatment has been shown to induce accumulation and extensive phosphorylation of p53 in epithelial tumor cells.^44^

**Figure 6.**
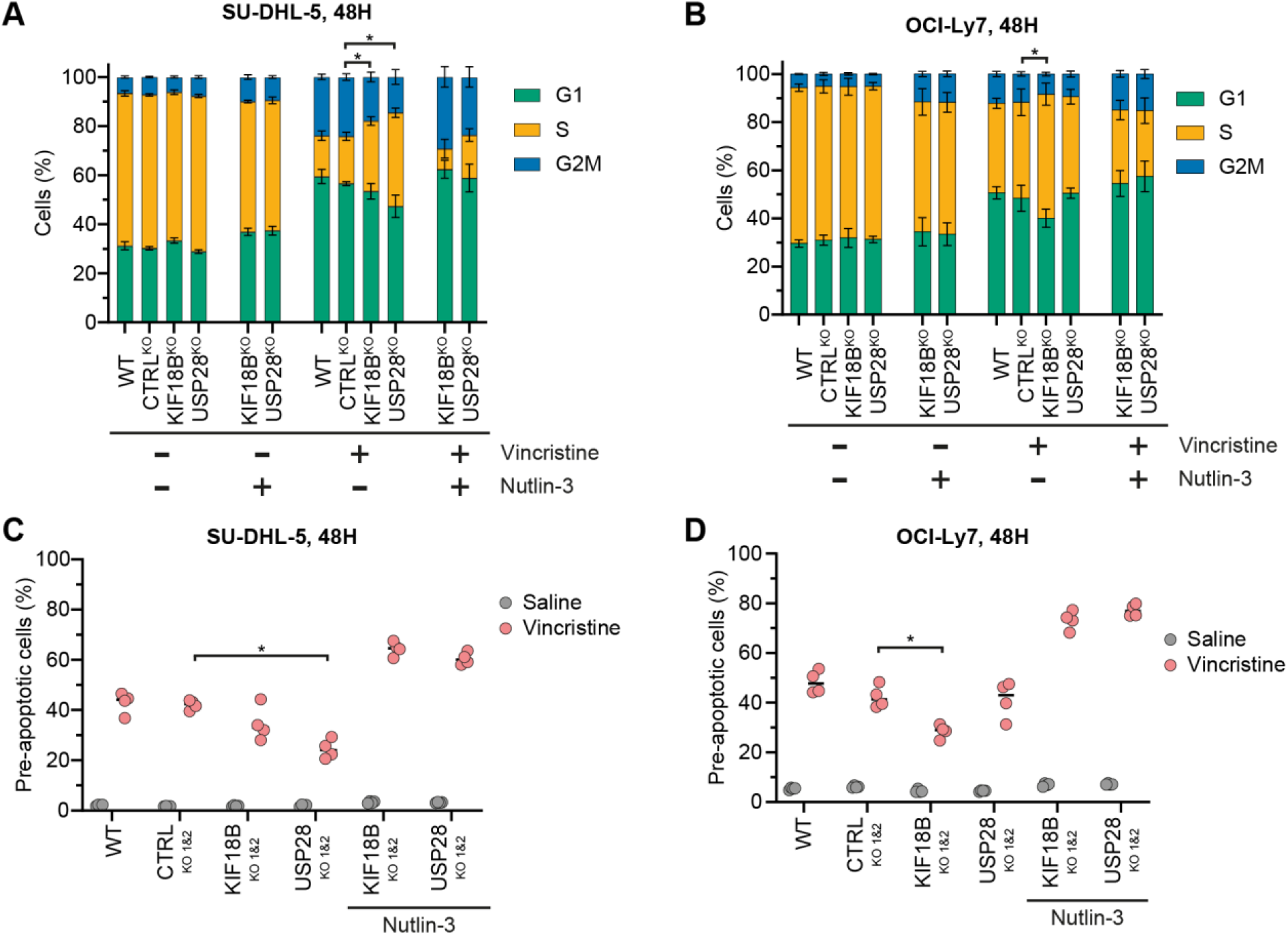
*KIF18B* and *USP28* knockout affects vincristine-induced cell cycle arrest and apoptosis. (**A-B**) Cell cycle analysis for SU-DHL-5 (**A**) and OCI-Ly7 (**B**) cells showing the percentage of cells in either G1-, S-, or G2/M-phase measured by incorporation of the nucleoside-analog EdU and DNA binding of propidium iodide by flow cytometry. (**C-D**) Percentage of pre-apoptotic cells in SU-DHL-5 (**C**) and OCI-Ly7 (**D**) measured by proteolytic activity of caspase-3 and caspase-7 in cleaving a fluorogenic substrate in live cells by flow cytometry. For SU-DHL-5, 2 ng/mL vincristine and 1.0 µg/mL nutlin-3 were used and for OCI-Ly7, 3.0 ng/mL vincristine and 2.5 µg/mL nutlin-3 were used.

Next, we examined the effect of *KIF18B* and *USP28* knockouts on vincristine-directed induction of apoptosis by measuring the proteolytic activity of caspase-3 and caspase-7. In SU-DHL-5 control samples, vincristine treatment induced 42% pre-apoptotic cells, which was reduced to 35% and 25% for *KIF18B* and *USP28* knockout samples, respectively (Figure 6C). In OCI-Ly7 control samples, vincristine treatment induced 42% pre-apoptotic cells, which was reduced to 29% for KIF18B knockout, whereas there was no change for USP28 knockout (Figure 6D). By treating the cells with nutlin-3, the reduction in apoptosis in the *KIF18B* and *USP28* knockout samples was reverted. This effect was only seen for samples treated with vincristine, whereas nutlin-3 had no effect on apoptosis in saline-treated samples, supporting that vincristine resulted in activation of p53 to induce apoptosis. Collectively, these results demonstrate that *KIF18B* and *USP28* are mediators of vincristine-induced cell cycle arrest and apoptosis.

## Discussion

The heterogeneity of DLBCL makes it challenging to predict the best treatment strategy, especially for high-risk patients.^2^ To address this issue, clinical trials have investigated ex vivo platforms for pharmacological testing as an approach to identifying personalized treatment options.^45,46^ We show that genome-wide CRISPR screening using a drug as selective pressure can be used to identify drug targets that result in substantial synergistic effects. CRISPR screening excels as a high-throughput method that enables simultaneous testing of thousands of drug targets using only a fraction of the cells required for testing single drugs one or two at a time. Our screening approach identified genes related to mitotic spindle organization as targets for sensitizing cells to vincristine treatment including KIF15 and KIF2A (Figure 7A).^47,48^ This was verified for the kinesin KIF18A using the inhibitor BTB-1. KIF18A mediates chromosome alignment during metaphase using its plus-end-directed motor on kinetochore microtubules and by regulating spindle length.^36^ Loss of KIF18A has been shown to result in high chromosome instability in aneuploid cells that showed particular dependence on KIF18A compared to near-diploid cancer cells during SAC inhibition treatment.^49^

**Figure 7.**
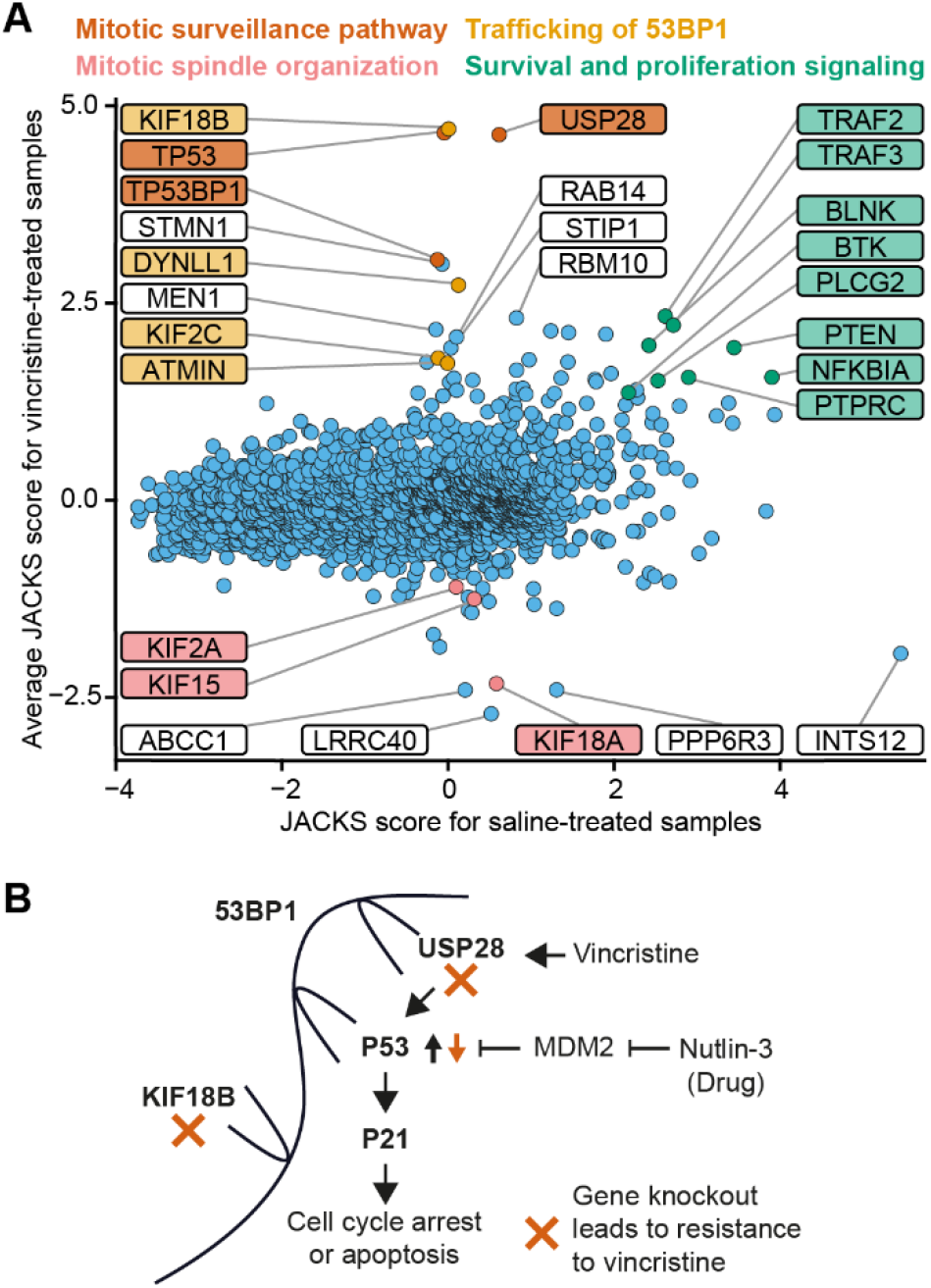
Inhibition of the mitotic surveillance pathway induces resistance to vincristine. (**A**) The average JACKS scores for the vincristine samples plotted against JACKS scores for the saline-treated samples. Color-highlighted genes are connected to the mechanism indicated in the same color above. (**B**) The proposed hypothesis. Vincristine induces the mitotic surveillance pathway entailing that the scaffolding protein 53BP1 binds USP28 and p53 enabling USP28 to de-ubiquitinate and thus, stabilize p53 from the ubiquitination from MDM2. The stabilized and activated p53 further activates p21 resulting in cell cycle arrest and apoptosis. KIF18B mediates this mechanism by binding and ensuring trafficking of 53BP1.

Our screening approach also identified genes related to BCR and TNFR signaling, which upon knockout increased the tolerance to vincristine and improved the survival and proliferation rate during normal cell culture (Figure 7A). BCR-signaling through BLNK, BTK, and PLCG2 follows the canonical NF-κB pathway causing the release and translocation of p65 (*RELA*), C-Rel (*REL*) and p50 (*NFKB1*) transcription factors, whereas TRAF2 and TRAF3 inhibit TNFR-signaling in the non-canionical NF-κB pathway that leads to translocation of Rel-B (*RELB*) and p52 (*NFKB2*).^35^ This suggests that for SU-DHL-5 cells activation of Rel-B and p52 is beneficial, whereas activation of p65, C-Rel and p50 is not. In fact, our screening data showed that *RELB* and *NFKB2* were essential for SU-DHL-5 cells with JACKS scores for saline-treated samples on -2.6 and -1.7, respectively, whereas *RELA, REL*, and *NFKB1* were not essential (scores on -0.1, 0.1, and -0.9, respectively). Previously, several studies have shown that GCB-subtype DLBCL cells rely primarily on BCR signaling through the PI3K-AKT-mTOR pathway, but do not depend on NF-κB signaling.^19,50,51^ However, in agreement with our results, a subset of GCB-subtype DLBCL cells have been shown to rely on NF-κB signaling through the Rel-B transcription factor.^52^

Recently, three studies used genome-wide CRISPR screening to uncover the cellular response to centrosome loss.^53–55^ All three studies identified USP28, p53 and 53BP1 as essential for inducing cell cycle arrest and apoptosis through p21 activation upon centrosome loss or prolonged mitosis. The three studies showed that USP28 and p53 bind to the scaffolding protein 53BP1, where USP28 de-ubiquitinates and, thus, stabilizes p53. They identified this response as a mitotic surveillance pathway independent of the SAC and the DNA damage checkpoint.^40^ In addition, another study using genome-wide CRISPR screening identified the mitotic surveillance pathway as the major driver behind resistance to the antimitotic drug TH588.^56^ Furthermore, in the study by Oshima et al investigating the vincristine response in ALL, the genes *TP53, TP53BP1, USP28*, and *KIF18B* ranked as number 1, 8, 15, and 368 as LOF resistance genes, respectively.^15^ Altogether, this suggests that USP28 and 53BP1-mediated stabilization of p53 is an important response to antimitotic treatments.

It remains unclear how the mitotic surveillance pathway is initiated. Luessing et al showed that KIF18B binds 53BP1 and is responsible for trafficking 53BP1 to the chromatin in response to DNA damage to mediate double-stranded DNA break repair.^57^ We speculate that the stabilization of activated p53 by binding 53BP1 and USP28 is enabled by the kinesin KIF18B ensuring the trafficking of 53BP1. Thus, knockout of either *TP53, TP53BP1, USP28*, or *KIF18B* disrupts the mitotic surveillance pathway by opposing the stabilization and trafficking of activated p53, resulting in less cell cycle arrest and apoptosis and, hence, increased tolerance to vincristine (Figure 7B). This would explain why the lack of *KIF18B*, but not of *USP28*, resulted in less cell cycle arrest and apoptosis for the TP53-mutated OCI-Ly7. As the amount of p53 protein already is high, USP28-mediated stabilization of p53 has little effect, whereas KIF18B-mediated trafficking of 53BP1 enables induction of cell cycle arrest and apoptosis by activated p53. In support of this, our screening data revealed multiple genes related to 53BP1-trafficking for which knockout induced vincristine resistance (Figure 7A). One of the top-ranked LOF genes inducing vincristine resistance was *DYNLL1*, a member of the cytoplasmic dynein 1 complex, which is involved in intracellular trafficking along microtubules. Interestingly, DYNLL1 also binds 53BP1,^58^ and another of the top-ranked genes *ATMIN* encodes a transcription factor responsible for *DYNLL1* gene transcription.^59^ Another top-ranked gene *KIF2C* encodes a kinesin-13 motor family member, which is known to be involved in trafficking along microtubules in DNA damage responses and has been shown to co-localize and co-migrate with 53BP1.^60^

A recent study showed that p53 transiently localizes to centrosomes during mitosis and that inhibition of this localization leads to the accumulation of activated p53 in foci that are required for recruitment of 53BP1 to mediate the stabilization of activated p53.^61^ Vincristine may interfere with the localization of p53 at centrosomes either directly by interacting with the microtubular structure at the centrosome or indirectly by inducing chromosome instability, leading to misregulated centrosomes. Interestingly, clinically relevant doses of the microtubule-binding drug paclitaxel primarily exerts a cytotoxic effect by inducing chromosome instability leading to multipolar spindles and misregulated centrosomes.^62^

It has long been suggested that a p53-dependent checkpoint controls correct mitosis in interphase.^63,64^ However, it has been unclear whether this checkpoint functions independently from the DNA damage checkpoint in interphase and from the intensification of apoptotic signaling during the SAC that pauses mitosis at metaphase.^8,65^ Notably, in our screening data knockout of genes responsible for the induction of the SAC or for recognizing DNA damage did not affect the vincristine response.

*TP53* is the most frequently mutated gene in cancer and is a recurrent mutation in relapsed/refractory DLBCL.^66^ In this study and in Laursen et al,^28^ the *TP53*-mutated cell line OCI-Ly7 was more resistant to vincristine than the *TP53* wildtype cell line SU-DHL-5. Our study suggests that the effect of vincristine in killing cancerous cells relies on a p53 response implicating that vincristine is less effective in loss-of-function *TP53*-mutated cells.

Based on genome-wide CRISPR screens, this work identifies new DLBCL drug targets, for which small molecular inhibitors may potentially support the effect of vincristine. We find that BTB-1, an inhibitor of KIF18A, in combination with vincristine shows synergistic effects in eradicating DLBCL cells. Through this work, we also demonstrate that dysregulation of the mitotic surveillance pathway unveils novel mechanisms supporting vincristine resistance in B-cells. Altogether, our findings contribute to a molecular understanding of resistance to vincristine in DLBCL, which may potentially support future drug development.

## Acknowledgements

The authors would like to thank FACS Core Facility, Aarhus University, Denmark for guidance and access to flow cytometers. This work was made possible through support from the following funding agencies: Independent Research Fund Denmark, The Carlsberg Foundation, NEYE-Fonden, Direktør Jacob Madsens og Olga Madsens Fond, Dagmar Marshalls Fond, Fabrikant Einar Willumsens Mindelegat, Raimond og Dagmar Ringgård-Bohns Fond, Andersen-Isted Fonden, Helga og Peter Kornings Fond.

## Authorship contributions

ABR, EAT, KD and JGM conceptualized the study. ABR and EAT performed the CRISPR screen. YL performed NGS. ABR performed bioinformatics analyses. ABR, IN and TWS performed functional studies. ABR and JGM wrote the manuscript. All authors approved the final manuscript.

## Disclosure of conflicts of interest

The authors declare no conflict of interest.

**Appendix**

Appendix 1: Supplementary figures to main article.

Appendix 2: Supplementary materials and methods, flow cytometry gating strategy, live cell count from drug assays, and uncropped western blots.

## Figure legends

**Supplementary Figure S1.**
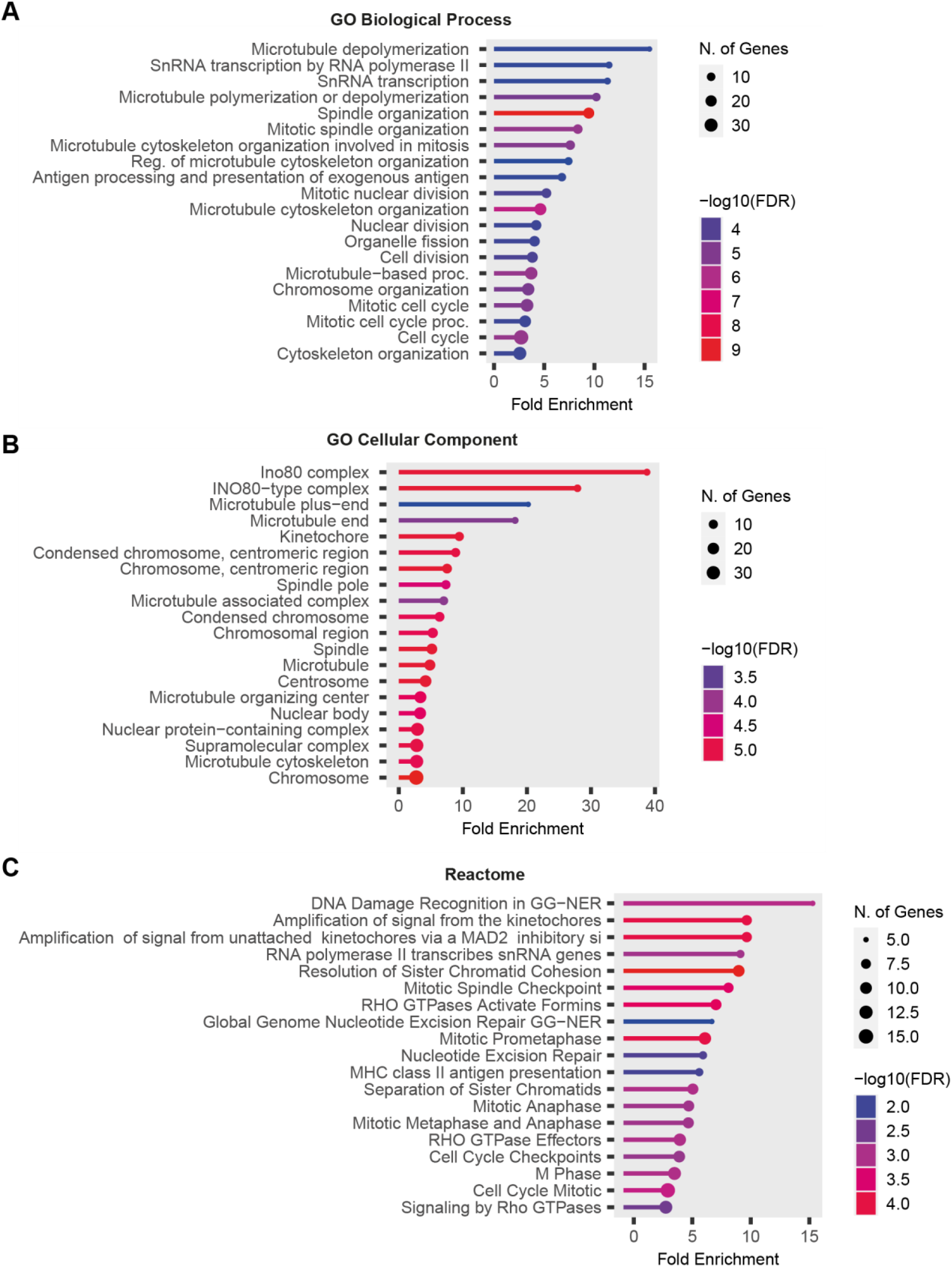
Gene-set enrichment analysis for the sensitizing LOF genes. (**A-C**) Performed with ShinyGO using the GO Biological Process (**A**), GO Cellular Component (**B**), and Reactome (**C**) gene sets. The color and size of the dots represent the false discovery rate (FDR) and the number of genes annotated to the specific pathway, respectively.

**Supplementary Figure S2.**
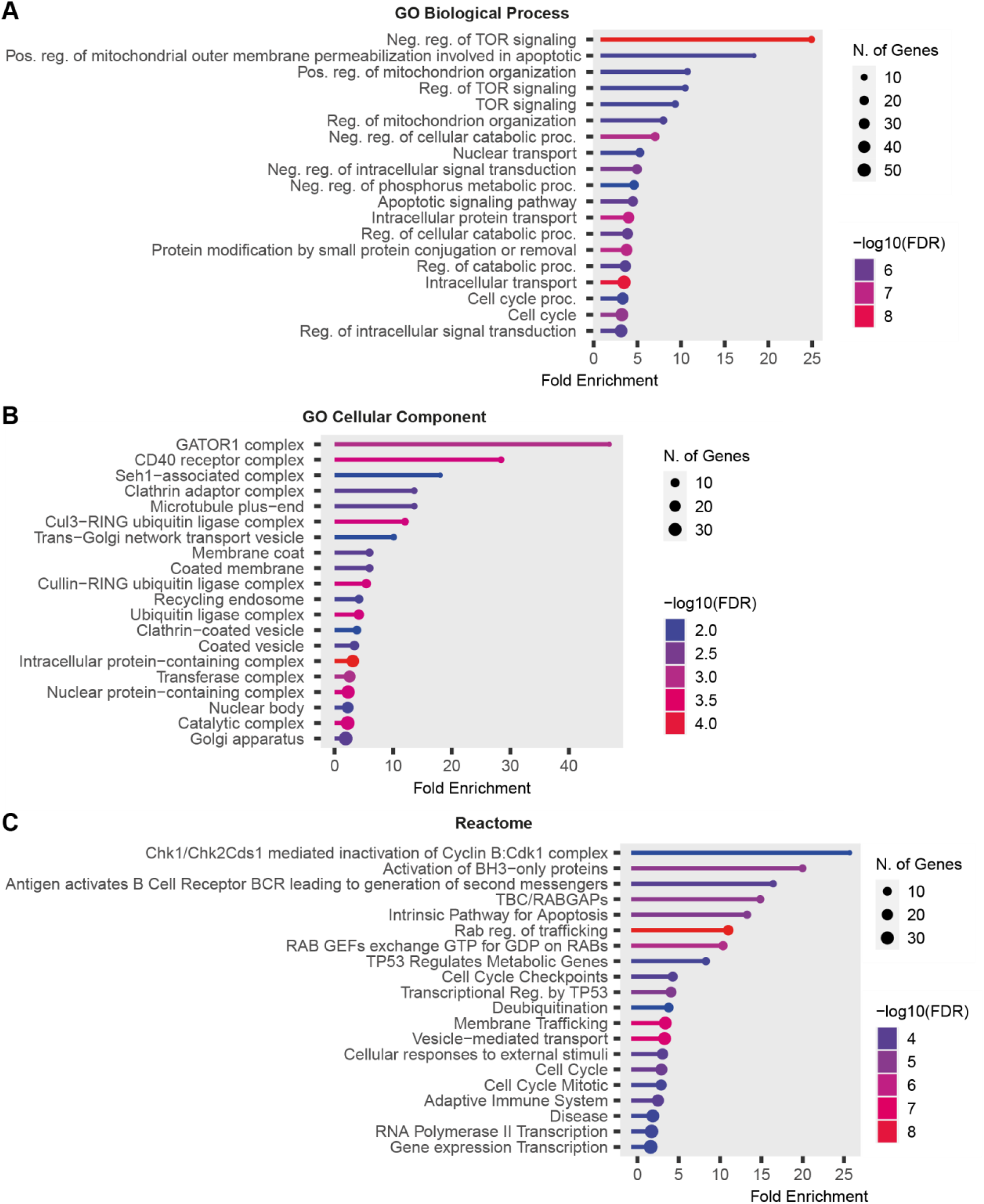
Gene-set enrichment analysis for the resistance-inducing LOF genes. (**A-C**) Performed with ShinyGO using the GO Biological Process (**A**), GO Cellular Component (**B**), and Reactome (**C**) gene sets. The color and size of the dots represent the false discover rate (FDR) and the number of genes annotated to the specific pathway, respectively.

**Supplementary Figure S3.**
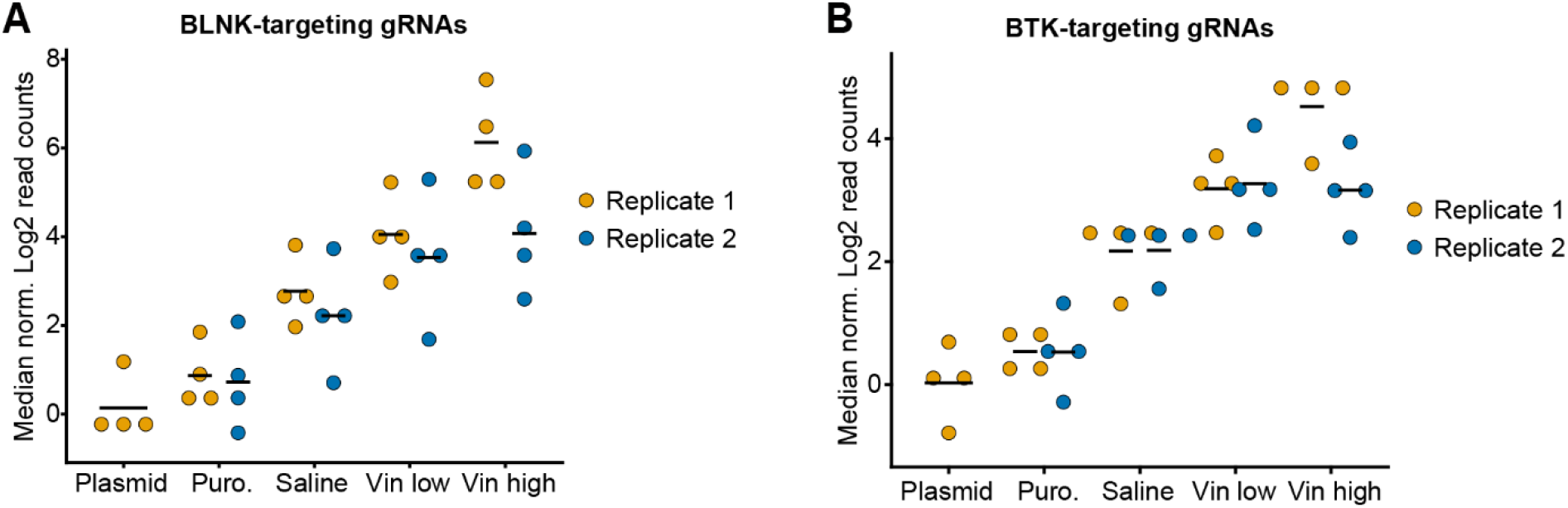
Distribution of gRNAs in genome-wide CRISPR screening. (**A-B**) Median normalized log2 read counts of gRNAs targeting *BLNK* (**A**) and *BTK* (**B**) across the different samples in the CRISPR screening.

**Supplementary Figure S4.**
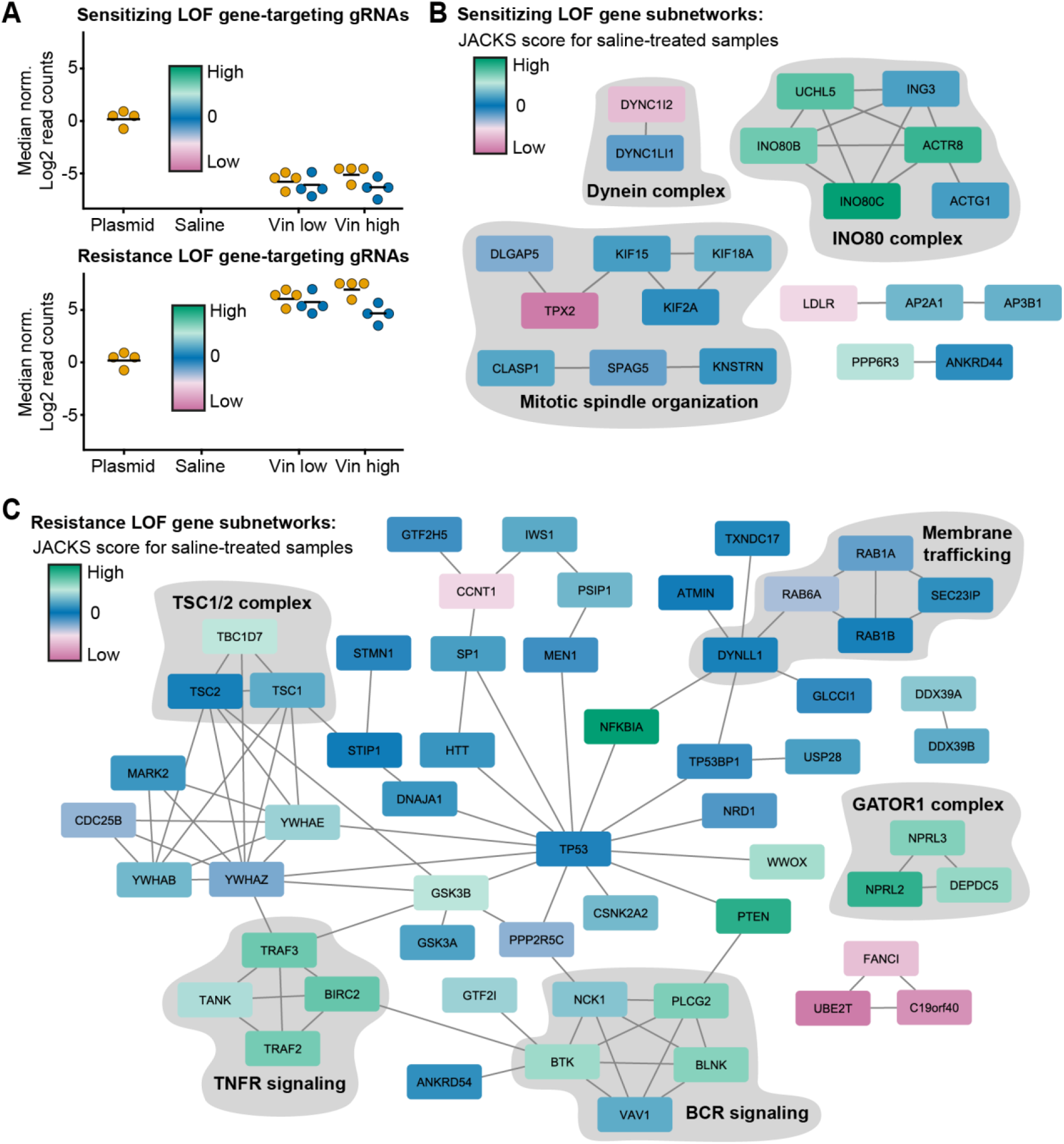
FDRnet-identified subnetworks of important genes color-coded according to their significance during normal cell culture. (**A**) The color code indicates the JACKS score for saline-treated samples using the plasmid sample as baseline. (**B-C**) The FDRnet-identified subnetworks among the most sensitizing (**B**) and resistance-inducing gene knockouts to vincristine treatment.

**Supplementary Figure S5.**
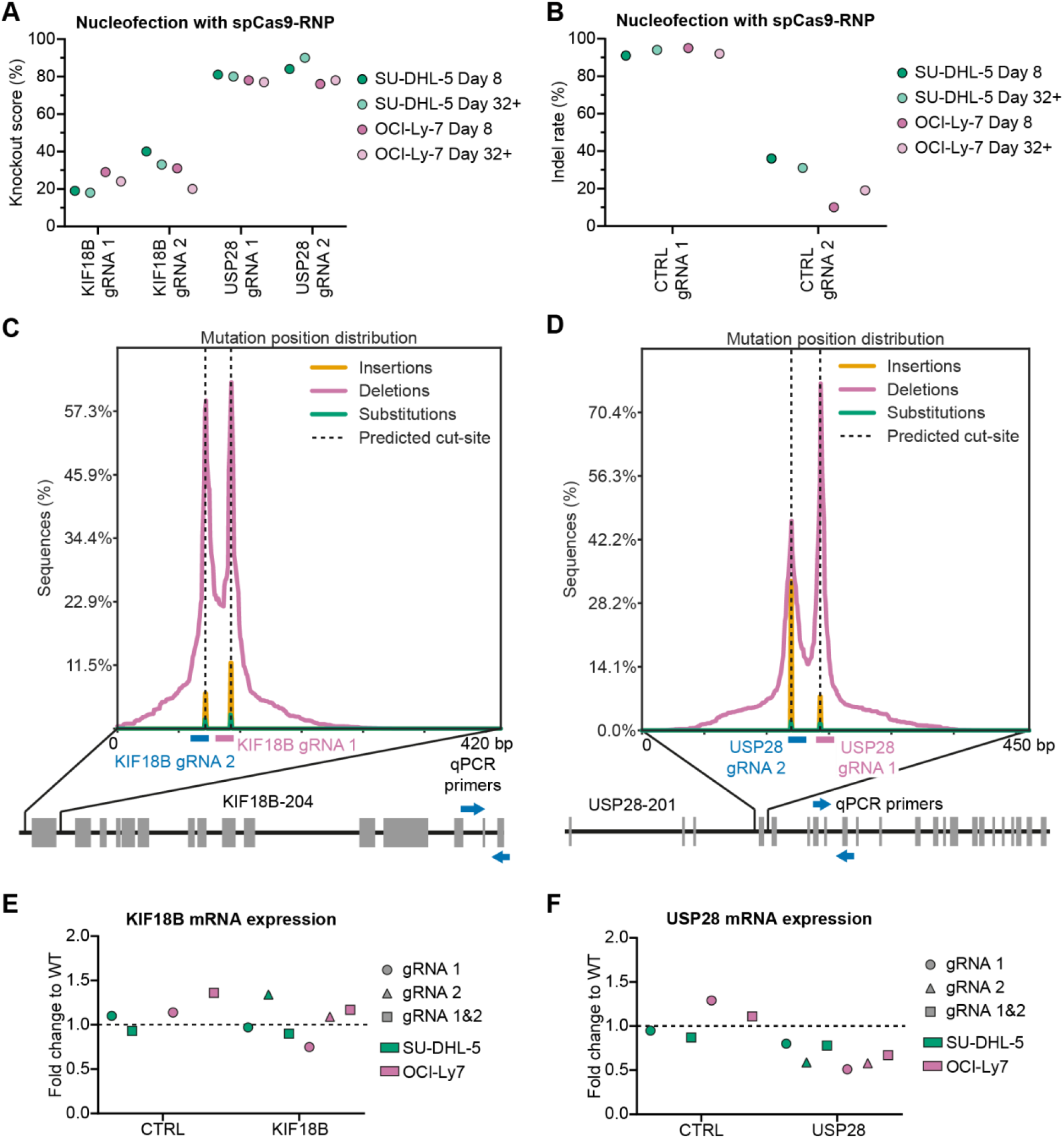
Generating *KIF18B* and *USP28* knockout cell lines. (**A**) Knockout rates measured after Cas9-induced gene knockout using Sanger sequencing at day 8 or more than 32 days after nucleofection. Knockout rates defined as the percentage of cells with either a frameshift mutation or indel larger than 21 bp. (**B**) Indel rates using control gRNAs targeting safe harbors. Measured after Cas9-induced gene knockout using Sanger sequencing at day 8 or more than 32 days after nucleofection. (**C-D**) Frequency of detected mutation patterns measured by NGS-mediated deep sequencing. Shown for the *KIF18B* (**C**) and *USP28* loci, where the primers used for qPCR are also indicated. Here represented by data for SU-DHL-5, which was comparable to data for OCI-L7. (**E-F**) mRNA expression of *KIF18B* (**E**) and *USP28* (**F**) in gene knockout-generated cell lines.

## Supplementary data

Supplementary table: Read counts of gRNAs and JACKS gene scores from genome-wide CRISPR screening.

## References

1. Howlader, N. et al. Cancer Statistics Review, 1975-2016 - SEER Statistics. Based on November 2018 SEER data submission, posted to the SEER web site, April 2019. Available at: https://seer.cancer.gov/csr/1975_2016/. (Accessed: 13th October 2021)

2. Hill, B. T. & Kahl, B. Upfront therapy for diffuse large B-cell lymphoma: looking beyond R-CHOP. https://doi.org/10.1080/17474086.2022.2124156 15, 805–812 (2022).

3. Tilly, H. et al. Polatuzumab Vedotin in Previously Untreated Diffuse Large B-Cell Lymphoma. N. Engl. J. Med. 386, 351–363 (2022).

4. Schmitz, R. et al. Genetics and Pathogenesis of Diffuse Large B-Cell Lymphoma. N. Engl. J. Med. 378, 1396–1407 (2018).

5. Chapuy, B. et al. Molecular subtypes of diffuse large B cell lymphoma are associated with distinct pathogenic mechanisms and outcomes. Nat. Med. 24, 679–690 (2018).

6. Lacy, S. E. et al. Targeted sequencing in DLBCL, molecular subtypes, and outcomes: a Haematological Malignancy Research Network report. Blood 135, 1759–1771 (2020).

7. DeVita, V. T. & Chu, E. A History of Cancer Chemotherapy. Cancer Res. 68, 8643– 8653 (2008).

8. Shi, J., Mitchison, T. J., Shi, J. & Mitchison, T. J. Cell death response to anti-mitotic drug treatment in cell culture, mouse tumor model and the clinic. Endocr. Relat. Cancer 24, T83–T96 (2017).

9. Matthews, H. K., Bertoli, C. & de Bruin, R. A. M. Cell cycle control in cancer. Nat. Rev. Mol. Cell Biol. 2021 231 23, 74–88 (2021).

10. Shalem, O. et al. Genome-scale CRISPR-Cas9 knockout screening in human cells. Science (80-.). 343, 84–87 (2014).

11. Wang, T., Wei, J. J., Sabatini, D. M. & Lander, E. S. Genetic screens in human cells using the CRISPR-Cas9 system. Science (80-.). 343, 80–84 (2014).

12. Koike-Yusa, H., Li, Y., Tan, E. P., Velasco-Herrera, M. D. C. & Yusa, K. Genome-wide recessive genetic screening in mammalian cells with a lentiviral CRISPR-guide RNA library. Nat. Biotechnol. 32, 267–273 (2014).

13. Thomsen, E. A. et al. Identification of BLNK and BTK as mediators of rituximab-induced programmed cell death by CRISPR screens in GCB-subtype diffuse large B-cell lymphoma. Mol. Oncol. 1878–0261.12753 (2020). doi:10.1002/1878-0261.12753

14. Palmer, A. C., Chidley, C. & Sorger, P. K. A curative combination cancer therapy achieves high fractional cell killing through low cross resistance and drug Additivity. Elife 8, (2019).

15. Oshima, K. et al. Mutational and functional genetics mapping of chemotherapy resistance mechanisms in relapsed acute lymphoblastic leukemia. Nat. Cancer 2020 111 1, 1113–1127 (2020).

16. Mo, Z. et al. Deciphering the mechanisms of CC-122 resistance in DLBCL via a genome-wide CRISPR screen. Blood Adv. 5, 2027–2039 (2021).

17. Tong, K. I. et al. Combined EZH2 Inhibition and IKAROS Degradation Leads to Enhanced Antitumor Activity in Diffuse Large B-cell Lymphoma. Clin. Cancer Res. 27, 5401–5414 (2021).

18. Reddy, A. et al. Genetic and Functional Drivers of Diffuse Large B Cell Lymphoma. Cell 171, 481–494.e15 (2017).

19. Phelan, J. D. et al. A multiprotein supercomplex controlling oncogenic signalling in lymphoma. Nature 560, 387–391 (2018).

20. Nie, M. et al. Genome-wide CRISPR screens reveal synthetic lethal interaction between CREBBP and EP300 in diffuse large B-cell lymphoma. Cell Death Dis. 12, 419 (2021).

21. Ryø, L. B., Thomsen, E. A. & Mikkelsen, J. G. Production and Validation of Lentiviral Vectors for CRISPR/Cas9 Delivery. Methods Mol. Biol. 1961, 93–109 (2019).

22. Thomsen, E. A. & Mikkelsen, J. G. CRISPR-Based Lentiviral Knockout Libraries for Functional Genomic Screening and Identification of Phenotype-Related Genes. in 343– 357 (Humana Press, New York, NY, 2019). doi:10.1007/978-1-4939-9170-9_21

23. Hart, T., Brown, K. R., Sircoulomb, F., Rottapel, R. & Moffat, J. Measuring error rates in genomic perturbation screens: gold standards for human functional genomics. Mol. Syst. Biol. 10, 733 (2014).

24. Allen, F. et al. JACKS: joint analysis of CRISPR/Cas9 knock-out screens.

25. Ge, S. X., Jung, D., Jung, D. & Yao, R. ShinyGO: a graphical gene-set enrichment tool for animals and plants. Bioinformatics 36, 2628–2629 (2020).

26. Yang, L., Chen, R., Goodison, S. & Sun, Y. An efficient and effective method to identify significantly perturbed subnetworks in cancer. Nat. Comput. Sci. 2021 11 1, 79–88 (2021).

27. Clement, K. et al. CRISPResso2 provides accurate and rapid genome editing sequence analysis. Nat. Biotechnol. 2019 373 37, 224–226 (2019).

28. Laursen, M. B. et al. Human B-cell cancer cell lines as a preclinical model for studies of drug effect in diffuse large B-cell lymphoma and multiple myeloma. Exp. Hematol. 42, 927–938 (2014).

29. Doench, J. G. et al. Optimized sgRNA design to maximize activity and minimize off-target effects of CRISPR-Cas9. Nat. Biotechnol. 34, 184–191 (2016).

30. Falgreen, S. et al. Predicting response to multidrug regimens in cancer patients using cell line experiments and regularised regression models. BMC Cancer 15, 1–15 (2015).

31. Li, W. et al. Quality control, modeling, and visualization of CRISPR screens with MAGeCK-VISPR. Genome Biol. 16, 1–13 (2015).

32. Cole, S. P. C. Targeting Multidrug Resistance Protein 1 (MRP1, ABCC1): Past, Present, and Future. http://dx.doi.org/10.1146/annurev-pharmtox-011613-135959 54, p95–117 (2014).

33. Eustermann, S. et al. Structural basis for ATP-dependent chromatin remodelling by the INO80 complex. Nat. 2018 5567701 556, 386–390 (2018).

34. Liu, G. Y. & Sabatini, D. M. mTOR at the nexus of nutrition, growth, ageing and disease. Nat. Rev. Mol. Cell Biol. 2020 214 21, 183–203 (2020).

35. Pasqualucci, L. & Klein, U. NF-κB Mutations in Germinal Center B-Cell Lymphomas: Relation to NF-κB Function in Normal B Cells. (2022). doi:10.3390/biomedicines10102450

36. Weaver, L. N. et al. Kif18A Uses a Microtubule Binding Site in the Tail for Plus-End Localization and Spindle Length Regulation. Curr. Biol. 21, 1500–1506 (2011).

37. Locke, J. et al. Structural basis of human kinesin-8 function and inhibition. Proc. Natl. Acad. Sci. U. S. A. 114, E9539–E9548 (2017).

38. McHugh, T., Gluszek, A. A. & Welburn, J. P. I. Microtubule end tethering of a processive kinesin-8 motor Kif18b is required for spindle positioning. J. Cell Biol. 217, 2403–2416 (2018).

39. Wang, X. et al. Targeting deubiquitinase USP28 for cancer therapy. Cell Death Dis. 2018 92 9, 1–10 (2018).

40. Lambrus, B. G. & Holland, A. J. A New Mode of Mitotic Surveillance. Trends Cell Biol. 27, 314–321 (2017).

41. Joerger, A. C., Ang, H. C. & Fersht, A. R. Structural basis for understanding oncogenic p53 mutations and designing rescue drugs. Proc. Natl. Acad. Sci. U. S. A. 103, 15056– 15061 (2006).

42. Lahav, G. et al. Dynamics of the p53-Mdm2 feedback loop in individual cells. Nat. Genet. 2004 362 36, 147–150 (2004).

43. Vassilev, L. T. et al. In Vivo Activation of the p53 Pathway by Small-Molecule Antagonists of MDM2. Science (80-.). 303, 844–848 (2004).

44. Stewart, Z. A., Luo Jia Tang & Pietenpol, J. A. Increased p53 phosphorylation after microtubule disruption is mediated in a microtubule inhibitor- and cell-specific manner. Oncogene 2001 201 20, 113–124 (2001).

45. Spinner, M. A. et al. Ex vivo drug screening defines novel drug sensitivity patterns for informing personalized therapy in myeloid neoplasms. Blood Adv. 4, 2768–2778 (2020).

46. Goh, J. et al. An ex vivo platform to guide drug combination treatment in relapsed/refractory lymphoma. Sci. Transl. Med. 14, eabn7824 (2022).

47. Sturgill, E. G. & Ohi, R. Kinesin-12 differentially affects spindle assembly depending on its microtubule substrate. Curr. Biol. 23, 1280–1290 (2013).

48. Uehara, R. et al. Aurora B and Kif2A control microtubule length for assembly of a functional central spindle during anaphase. J. Cell Biol. 202, 623–636 (2013).

49. Cohen-Sharir, Y. et al. Aneuploidy renders cancer cells vulnerable to mitotic checkpoint inhibition. Nat. 2021 5907846 590, 486–491 (2021).

50. Eric Davis, R., Brown, K. D., Siebenlist, U. & Staudt, L. M. Constitutive Nuclear Factor κB Activity Is Required for Survival of Activated B Cell–like Diffuse Large B Cell Lymphoma Cells. J. Exp. Med. 194, 1861–1874 (2001).

51. Havranek, O. et al. Tonic B-cell receptor signaling in diffuse large B-cell lymphoma. Blood 130, 995–1006 (2017).

52. Eluard, B. et al. The alternative RelB NF-κB subunit is a novel critical player in diffuse large B-cell lymphoma. Blood 139, 384–398 (2022).

53. Lambrus, B. G. et al. A USP28–53BP1–p53–p21 signaling axis arrests growth after centrosome loss or prolonged mitosis. J. Cell Biol. 214, 143–153 (2016).

54. Fong, C. S. et al. 53BP1 and USP28 mediate p53-dependent cell cycle arrest in response to centrosome loss and prolonged mitosis. Elife 5, (2016).

55. Meitinger, F. et al. 53BP1 and USP28 mediate p53 activation and G1 arrest after centrosome loss or extended mitotic duration. J. Cell Biol. 214, 155 (2016).

56. Gul, N. et al. The MTH1 inhibitor TH588 is a microtubule-modulating agent that eliminates cancer cells by activating the mitotic surveillance pathway. Sci. Reports 2019 91 9, 1–12 (2019).

57. Luessing, J. et al. The nuclear kinesin KIF18B promotes 53BP1-mediated DNA double-strand break repair. Cell Rep. 35, 109306 (2021).

58. Becker, J. R. et al. The ASCIZ-DYNLL1 axis promotes 53BP1-dependent non-homologous end joining and PARP inhibitor sensitivity. Nat. Commun. 2018 91 9, 1– 12 (2018).

59. Jurado, S. et al. ATM Substrate Chk2-interacting Zn2+ Finger (ASCIZ) Is a Bi-functional Transcriptional Activator and Feedback Sensor in the Regulation of Dynein Light Chain (DYNLL1) Expression. J. Biol. Chem. 287, 3156–3164 (2012).

60. Zhu, S. et al. Kinesin Kif2C in regulation of DNA double strand break dynamics and repair. Elife 9, (2020).

61. Contadini, C. et al. p53 mitotic centrosome localization preserves centrosome integrity and works as sensor for the mitotic surveillance pathway. Cell Death Dis. 2019 1011 10, 1–16 (2019).

62. Zasadil, L. M. et al. Cytotoxicity of paclitaxel in breast cancer is due to chromosome missegregation on multipolar spindles. Sci. Transl. Med. 6, (2014).

63. Cross, S. M. et al. A p53-Dependent Mouse Spindle Checkpoint. Science (80-.). 267, 1353–1356 (1995).

64. Lanni, J. S. & Jacks, T. Characterization of the p53-Dependent Postmitotic Checkpoint following Spindle Disruption. Mol. Cell. Biol. 18, 1055–1064 (1998).

65. Gascoigne, K. E. & Taylor, S. S. Cancer Cells Display Profound Intra- and Interline Variation following Prolonged Exposure to Antimitotic Drugs. Cancer Cell 14, 111– 122 (2008).

66. Morin, R. D. et al. Genetic Landscapes of Relapsed and Refractory Diffuse Large B-Cell Lymphomas. Clin. Cancer Res. 22, 2290–2300 (2016).

